# Evaluating the sensitivity of functional connectivity measures to motion artifact in resting-state fMRI data

**DOI:** 10.1101/2020.05.04.072868

**Authors:** Arun S. Mahadevan, Ursula A. Tooley, Maxwell A. Bertolero, Allyson P. Mackey, Danielle S. Bassett

## Abstract

Functional connectivity (FC) networks are typically inferred from resting-state fMRI data using the Pearson correlation between BOLD time series from pairs of brain regions. However, alternative methods of estimating functional connectivity have not been systematically tested for their sensitivity or robustness to head motion artifact. Here, we evaluate the sensitivity of six different functional connectivity measures to motion artifact using resting-state data from the Human Connectome Project. We report that FC estimated using full correlation has a relatively high residual distance-dependent relationship with motion compared to partial correlation, coherence and information theory-based measures, even after implementing rigorous methods for motion artifact mitigation. This disadvantage of full correlation, however, may be offset by higher test-retest reliability and system identifiability. FC estimated by partial correlation offers the best of both worlds, with low sensitivity to motion artifact and intermediate system identifiability, with the caveat of low test-retest reliability. We highlight spatial differences in the sub-networks affected by motion with different FC metrics. Further, we report that intra-network edges in the default mode and retrosplenial temporal sub-networks are highly correlated with motion in all FC methods. Our findings indicate that the method of estimating functional connectivity is an important consideration in resting-state fMRI studies and must be chosen carefully based on the parameters of the study.

## Introduction

Ever since the initial observation of correlations in spontaneous functional magnetic resonance imaging (fMRI) blood-oxygen level dependent (BOLD) signals acquired from subjects at rest, the field of resting-state functional connectivity has grown exponentially^1^. Functional connectivity has been used as a tool to explore large-scale features of human brain organization^2–6^, how this organization changes over the course of development^7–13^, and the association of such organization with individual behavior^14,15^. However, head motion artifact is a pervasive problem in functional connectivity analysis, decreasing certainty of findings and impacting subsequent interpretations. In-scanner head movements result in structured noise that leads to the spurious identification of putative functional connections, a problem further compounded by the fact that some individuals move systematically more than others^16^. Consequently, a number of research groups have developed statistical preprocessing methods to mitigate the impact of motion artifact; the development of such methods is an ongoing and active field of research in its own right^17–20^.

While much effort has been directed toward developing effective denoising pipelines to mitigate motion artifact, the subsequent estimation of functional connectivity in fMRI data has remained fairly constant. Functional connectivity (FC) between brain regions is typically estimated through a Pearson correlation between the BOLD time series of two regions of interest (ROIs). However, functional connectivity is formally defined as any statistical relation between time series^21^ and there exist many other statistical methods to compute similarity between time series. A few examples include coherence methods^22–24^, which compute similarity in frequency space, and methods based on information theory^25–27^, which quantify the amount of shared information between signals.

The performance of different FC estimation methods has been evaluated using generative models for a variety of neurophysiological data, including simulated BOLD signals^28,29^. These studies have typically focused on the ability of FC estimation methods to recover the underlying network structure from simulated BOLD data. Major findings include that full or partial correlation successfully recovers the underlying network structure in simulated data^28^. It is perhaps due to these results, and the associated ease of implementation, that correlation-based methods are so popular in the field. Yet, very few studies have used real fMRI data to compare the differential sensitivity or robustness of different FC estimation methods to motion artifact. The field awaits an appraisal of different FC estimation approaches with regard to their ability to overcome the specific type of noise introduced by motion artifact in fMRI data^16,30^.

In the present study, we used resting state fMRI data from the Human Connectome Project (HCP) to evaluate six different FC estimation methods: Pearson correlation, partial correlation, coherence, wavelet coherence, mutual information in the time domain, and mutual information in the frequency domain. The sensitivity of each of these methods to subject motion and their success in identifying network structure was evaluated using four benchmarks: (a) correlations of subject motion with edge weights after denoising (QC-FC correlations), (b) the distance-dependence of QC-FC correlations, (c) the degree to which canonical brain systems could be identified through modularity maximization, and (d) the extent to which the functional connectivity estimates could be reproduced in successive scans (test-retest reliability). Collectively, these efforts serve to inform our usage of FC estimation methods, and their relative strengths and weaknesses.

## Methods

In order to evaluate the differential sensitivity of different FC estimation methods to motion, we first applied common denoising pipelines to a large resting state dataset, estimated functional connectivity matrices using 6 different methods, and finally compared the performance of each of these estimates using a set of common quality control (QC) measures. Details of data preprocessing, FC estimation, and QC measures are described below.

### Data and preprocessing

In this study, we leveraged data from the S1200 release of the Human Connectome Project (HCP)^31^, a multi-site consortium that collected extensive MRI, behavioral, and demographic data from a large cohort of over 1000 subjects. As part of the HCP protocol, subjects underwent four separate resting-state scans, which included both left-right (REST1_LR, REST2_LR) and right-left (REST1_RL, REST2_RL) phase encoding directions. All functional connectivity data analyzed in this report came from these scans.

We used the ICA-FIX resting-state data provided by the Human Connectome Project, which used 24-parameter regression followed by ICA+FIX denoising to remove nuisance and motion signals^32,33^. In addition, we removed the mean global signal and bandpass filtered the time series from 0.009 to 0.08 Hz. All results reported in the main text were obtained with GSR and bandpass filtering applied, but we report results without application of GSR and bandpass filtering in the supplementary information. Further, we did not analyze subjects for whom greater than 50% of frames had a framewise displacement above 0.2 millimeters or a derivative root mean square above 75, leaving 778 subjects from REST1_LR, 800 from REST1_RL, 776 from REST2_LR, and 763 from REST2_RL. This threshold was chosen because it is typical for analyses of functional connectivity, and we wanted our conclusions about motion and functional connectivity to apply to common analysis pipelines^34–37^.

For each scan, we used the mean relative RMS (root-mean squared) displacement during realignment using MCFLIRT, provided by the Human Connectome Project, as our primary measure of motion. Summary statistics of the cohorts analyzed and their head motion are shown in Table 1.

**Table 1.** Summary statistics of motion in a cohort taken from the HCP S1200 release

| Scan | REST1_LR | REST1_RL | REST2_LR | REST2_RL |
| --- | --- | --- | --- | --- |
| Number of subjects analyzed (n) | 778 | 800 | 776 | 763 |
| Mean relative RMS motion (average) | 0.079 | 0.077 | 0.078 | 0.079 |
| Mean relative RMS motion (standard deviation) | 0.022 | 0.021 | 0.021 | 0.023 |

From the preprocessed data, we estimated mean BOLD time series using two cortical parcellation schemes: the 333-node Gordon parcellation^38^ and the 100-node Schaefer parcellation^35^. For all scans, the MSMAII registration was used, and the mean time series of vertices on the cortical surface (fsL32K) in each parcel was calculated.

### Description of functional connectivity measures

In the present study, we evaluated 6 different methods for estimating functional connectivity from BOLD time series data. Here we provide a brief overview of the methods evaluated.

### Pearson’s correlation coefficient

The Pearson correlation coefficient is a simple and commonly applied method to evaluate linear correlation between two time series. It is defined as the covariance between the two signals over time divided by the product of their standard deviations. Formally, the zero-order (without lag) Pearson correlation, *ρ*_*i,j*_, between the signals of regions *i* and *j* is given by

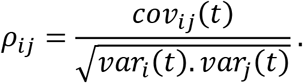

Note that *ρ*_*i,j*_ varies in the interval [−1, 1] with positive values indicating positive correlation and negative values indicating negative correlation.

### Partial correlation coefficient

Partial correlation is a measure of the linear correlation between two time series after regressing out the time series of all other nodes in the network. Partial correlation has been proposed as an effective method to distinguish between direct and indirect links between nodes^28^. Note that partial correlation also varies between [−1, 1] with positive values indicating positive correlation and negative values indicating negative correlation, after accounting for all other time series in the network.

### Mutual information (time domain)

The mutual information is a statistical measure of the shared information between two time series. The information content of a given time series *X*(*t*) can be defined through its Shannon entropy^25,26^, which is given by

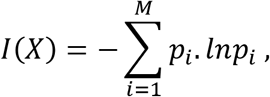

where *X*(*t*) is partitioned into *M* bins, with *p*_*i*_ representing the probability of the *i*-th bin. Now, the joint entropy between *X*(*t*) and a second time series *Y*(*t*) is defined as

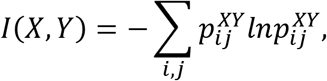

where 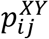 is the joint probability of *X* = *X*_*i*_ and *Y* = *Y*_*i*_. The mutual information between *X* and *Y* is then,

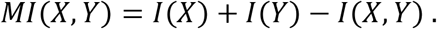

In order to obtain values in the range [0, 1], we computed the normalized mutual information^39^ as

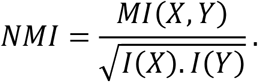

Thus, the normalized mutual information between two independent signals is 0 and has a maximum of 1 for identical signals. While correlation coefficients measure linear relationships, the mutual information is a statistical measure of both linear and non-linear relationships between time series.

### Coherence

Coherence is a measure of the cross-correlation between two signals in the frequency domain. At a given frequency *λ*, the coherence between the signal of region *i* and the signal of region *j* is given by

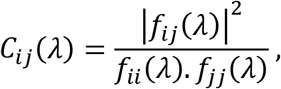

where *f*_*i,j*_(*λ*) is the cross-spectral density between signals *i* and *j*, and *f*_*ii*_ (*λ*) and *f*_*jj*_ (*λ*) are the auto-spectral densities of signal *i* and *j*, respectively. Note that *C*_*ij*_ (*λ*) varies in the interval [0, 1].

We evaluated coherence using the MATLAB toolbox for functional connectivity^40^, in which spectral densities are calculated using Welch’s averaged, modified periodogram method.

### Wavelet coherence

Wavelet coherence is a measure of the correlation between two signals in the time-frequency space. It is calculated in a similar manner to coherence, but spectral densities are calculated by convolving time series with wavelet functions such as the Morlet wavelet function^41^, which expand the signal in time-frequency space. We evaluated wavelet coherence using the Grinsted toolbox^42^.

### Mutual information (frequency domain)

Mutual information can also be evaluated based on coherence in the frequency domain^27^, defined for a given frequency range [*λ*1, *λ*2] as

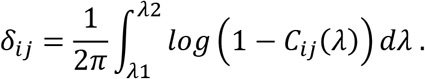

With a simple transformation, a normalized mutual information in the range [0, 1] can be obtained as

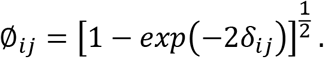

We used the implementation provided in the MATLAB toolbox for functional connectivity^40^.

### Overview of functional connectivity estimation

Using the 6 measures described above, we estimated functional connectivity between each pair of preprocessed BOLD times series, resulting in *n* × *n* matrices for each subject, where *n* is the number of parcels in the parcellation scheme. The estimated networks provide a description of interactions (edges) among brain regions (nodes) that could then be probed for various features of interest using network science^43^.

The 6 metrics evaluated can be broadly classified into categories based on their mode of operation. Pearson correlation, partial correlation, and mutual information (time) work in the time domain, whereas coherence and mutual information (frequency) work in the frequency domain, and wavelet coherence works in time-frequency space. For all the frequency-based methods, we evaluated the average connectivity in the frequency range [0.009Hz, 0.08Hz], which is the same frequency range at which all resting-state scans were bandpass filtered. Pearson correlation and partial correlation fall in the interval [−1, 1], and all the other metrics fall in the interval [0, 1] (Table 2).

**Table 2.** Overview of functional connectivity estimation measures, their range and domain of operation

| Domain | Metric | Range |
| --- | --- | --- |
| Time | Pearson correlation | [-1,1] |
|  | Partial correlation | [-1,1] |
|  | Mutual Information (time) | [0,1] |
| Frequency | Coherence | [0,1] |
|  | Mutual Information (frequency) | [0,1] |
| Time-frequency | Wavelet Coherence | [0,1] |

Correlation matrices are typically subjected to Fisher’s r-to-z transformation to normalize the range of values. Figure S2 shows that results do not change significantly when the transform is applied to the correlation matrices. Therefore, we report results in the main text without performing Fisher transforms on any of the matrices, in order to facilitate a more direct comparison between methods.

### Overview of outcome measures

We evaluated the sensitivity of each FC metric to subject motion using four benchmarks: residual QC-FC correlations, distance-dependence of QC-FC correlations, test-retest reliability, and the modularity quality index.

### Residual QC-FC correlations

Quality control-functional connectivity (QC-FC) correlations are a widely used benchmark to evaluate the efficacy of denoising pipelines applied in resting-state fMRI connectivity analysis^17,18^. Here we used this benchmark to evaluate residual motion artifact for each of the functional connectivity estimates after application of a common denoising pipeline. First, we computed functional connectivity using the 6 metrics described in the previous section, for the 333-node Gordon and 100-node Yeo parcellation schemes. We then computed the partial correlation between functional connectivity estimates for each edge and the relative mean RMS motion of each subject, controlling for subject age and sex, thus obtaining a distribution of edge-specific correlations with subject motion. From this distribution, we computed the percentage of edges for which the QC-FC correlations were statistically significant (p<0.05, no correction for multiple comparisons).

### Distance-dependence of QC-FC correlations

Motion artifact has been known to have a distance-dependent effect on FC estimates, thereby inflating the estimated strength of short-distance connections and reducing the estimated strength of long-distance connections^16,44^. To quantify this effect, we measured the correlation between the absolute values of QC-FC correlations (see above section) and the Euclidean distance separating the centroids of the node pair associated with each edge. This correlation served as a benchmark for the distance-dependence of the residual motion artifact.

### Test-retest reliability

To evaluate the reliability of functional connectivity estimates, we computed the intra-class correlation (ICC) across different resting-state scans performed on the same subject in the HCP dataset. The intra-class correlation coefficient *ρ* is defined^45^ as

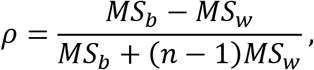

where *MS*_*b*_ is the between-subject mean squared strength of each edge, *MS*_*w*_ is the within-subject mean squared strength of each edge, and *n* is the number of scans per subject, which in this case is 4.

### System identifiability

To evaluate the possibility that more motion-resilient FC metrics might enable better detection of signals of interest, we consider the outcome measure of system identifiability^17,46,47^. We use the term *system* to refer to a set of brain regions that are strongly functionally connected; and we use the phrase *system identifiability* to refer to the ease with which such systems can be detected from functional connectivity matrices. We employed the modularity quality index, *Q*, as a measure of system identifiability. The modularity quality index is a quantification of the extent to which a network can be subdivided into groups or modules characterized by strong intramodular connectivity and weak intermodular connectivity. Such modularity is indicative of the assortative community structure commonly observed in functional brain networks^48,49^.

We estimated the modularity quality index for each subject’s network by maximizing the modularity quality function originally defined by Newman^46^ and subsequently extended to weighted and signed networks by Rubinov and Sporns^50^, among others^51,52^. For FC metrics resulting in weights falling within the interval [0,1], or results estimated from the absolute value of edge weights, we employed the weighted generalization of the modularity quality index. We first let the weight of a positive edge between nodes *i* and *j* be given by 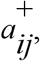, and the strength of a node *i*,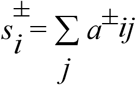, be given by the sum of the positive edge weights of *i*. We denote the chance expected within-module edge weights as 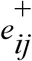 for positive weights where 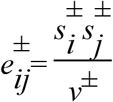. We let the total weight, 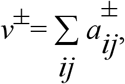, be the sum of all positive edge weights in the network. Then the weighted generalization of the modularity quality index is given by

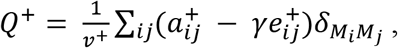

where *M_i_* is the community to which node *i* is assigned, and *M_j_* is the community to which node *j* is assigned. The Kronecker delta function, 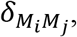, takes on a value of 1 when *M*_*i*_ = *M*_*j*_ and a value of 0 when *M*_*i*_ ≠ *M*_*j*_. The tunable structural resolution parameter, γ, scales the relative importance of the expected within-module weights (the null model) and in practice, the size of the communities; smaller or larger values of γ result in correspondingly larger or smaller communities. We use a Louvain-like locally greedy algorithm^53^ as a heuristic to maximize this modularity quality index subject to a partition *M* of nodes into communities.

For FC metrics resulting in weights falling in the interval [−1,1], specifically the Pearson and partial correlations, we employed the asymmetrically weighted generalization of *Q* suitable for networks containing negative weights^50^. Specifically, we follow Rubinov and Sporns^50^ by first letting the weight of a positive edge between nodes *i* and *j* be given by 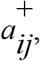, the weight of a negative edge between nodes *i* and *j* be given by 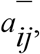, and the strength of a node *i*, 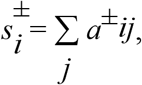 be given by the sum of the positive or negative edge weights of *i*. We denote the chance expected within-module edge weights as 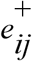 for positive weights and 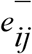 for negative weights, where 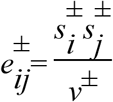. We let the total weight, 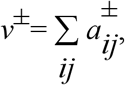, be the sum of all positive or negative edge weights in the network. Then an asymmetric signed generalization of the modularity quality index can be written as

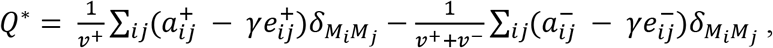

where *M_i_*, *M_j_*, 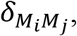, and γ are defined as above.

We examined average *Q* for each FC metric as a measure of system identifiability, as well as the partial correlation between *Q* and mean relative RMS for each subject while controlling for average network weight, age, and sex. Additionally, we addressed two potential confounds that have not been previously addressed in work examining *Q* as a measure of system identifiability: the number of communities *k* detected during modularity maximization, and the mean and distribution of edge weights in a given network (see Supplementary Methods for details).

## Results

### Characteristics of FC matrices computed using different methods

We first characterized the functional connectivity edge weights estimated using different methods. Figure 1 shows pairwise scatterplots between edge weights computed using all 6 methods. These plots show the non-linear relationships between edges estimated using Pearson correlation and non-correlation based methods. Of particular interest is the mapping from negative edges in Pearson correlation to others. For instance, the weights of negative edges in Pearson matrices have an inverse relationship with the weights of edges in wavelet coherence matrices – the more negative a Pearson edge weight, the higher its wavelet coherence edge weight.

**Figure 1.**
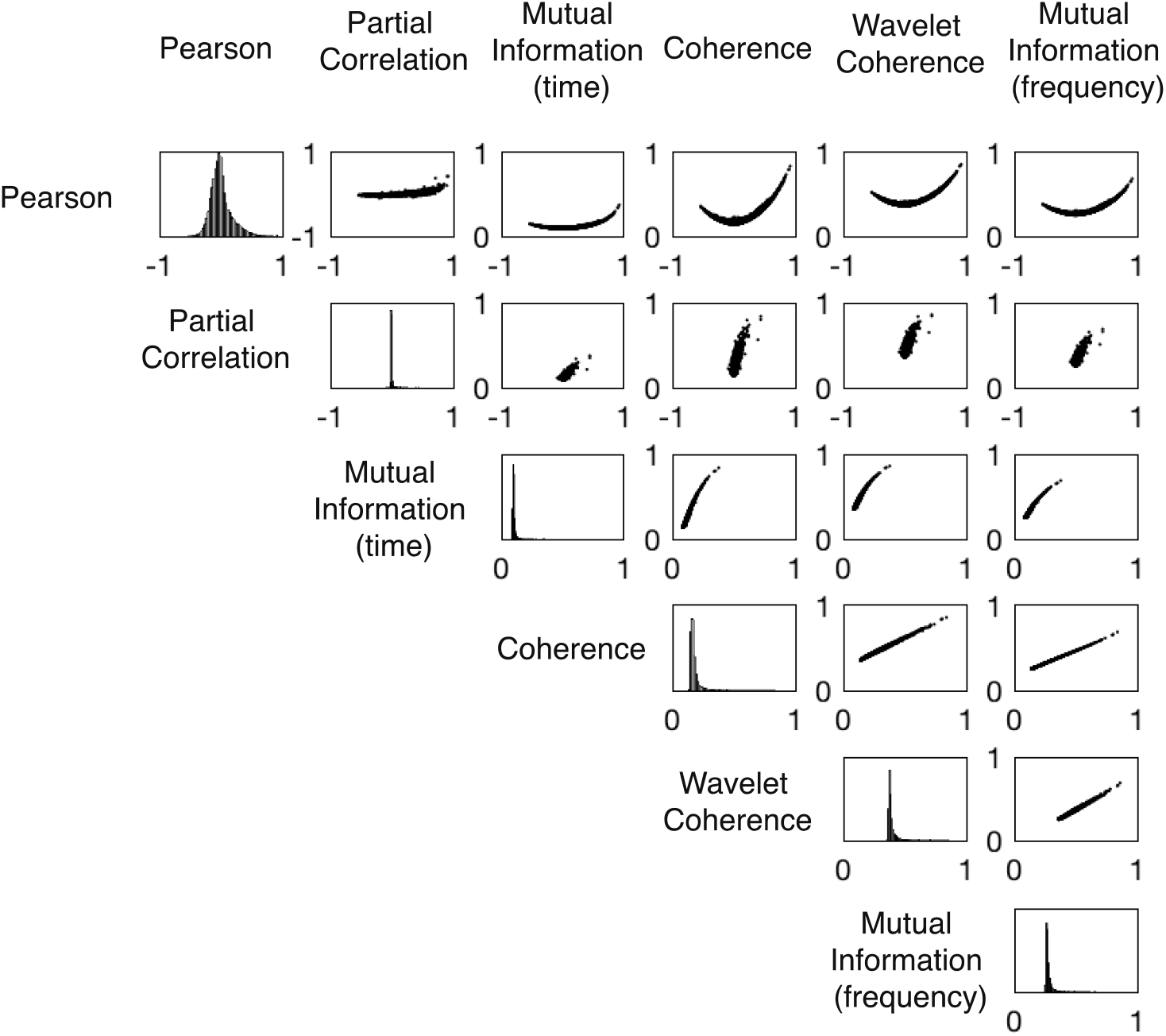
Edge weight correlations for different FC metrics. Pairwise scatter plots between edge weights calculated using different FC methods. Diagonal entries show histograms of edge weights. Results are shown for the REST1_LR scan, with FC estimated using the 333-node Gordon parcellation.

In Figure 2, FC matrices estimated using different methods are displayed as heatmaps, with canonical systems in the Gordon parcellation highlighted in the x and y color bars. Modular structure can clearly be seen in all matrices, with clean delineation of canonical systems in all matrices. Further, in Pearson correlation matrices, and to a lesser extent in partial correlation matrices, well known negative associations are apparent, for instance between the default mode and dorsal attention systems^54^.

**Figure 2.**
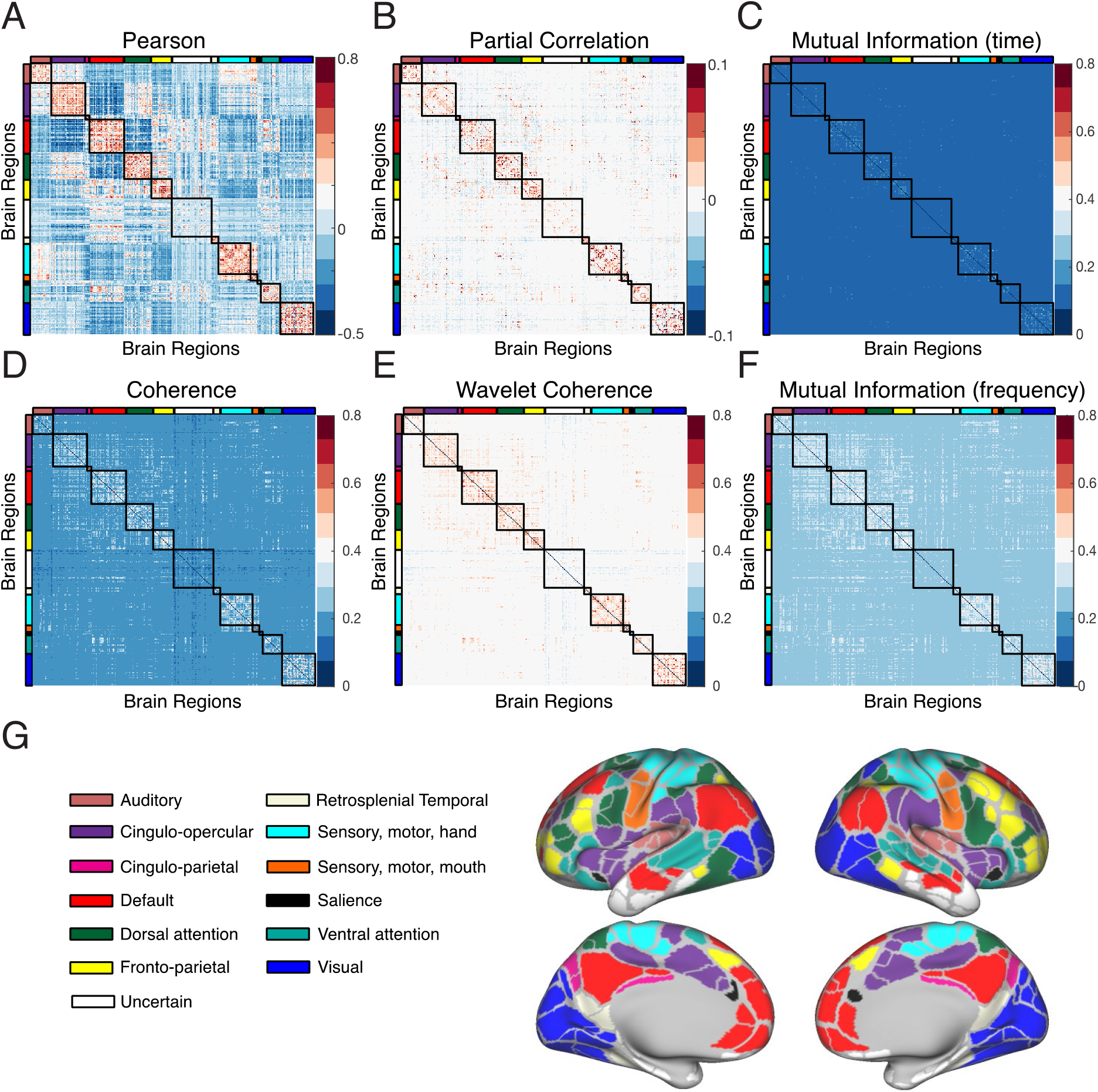
FC matrices using different estimation methods. Edge weight heatmaps are shown for the 333-node Gordon parcellation, with canonical systems labeled as colored bars. Each entry in the heatmap is the average edge weight across 778 subjects in the REST1_LR scans, estimated using **(A)** Pearson correlation, **(B)** partial correlation, **(C)** mutual information (time), **(D)** coherence, **(E)** wavelet coherence, and **(F)** mutual information (frequency). **(G)** Canonical systems for the Gordon parcellation displayed on the HCP S1200 group average cortical surface.

### Pearson correlation shows high residual QC-FC correlations

Next, we evaluated the sensitivity of edge weights (computed using different methods) to subject motion. We used the residual QC-FC correlation benchmark, which measures the edgewise relationship between the relative mean RMS motion of each subject and their estimated edge weights, after the application of denoising pipelines.

Our findings are summarized in Figure 3. Specifically, panels A and B of Figure 3 show the fraction of edges for which the QC-FC correlations are statistically significant (p<0.05, no correction for multiple comparisons). Significantly, partial correlation is the best-performing FC method, Pearson correlation is the worst-performing FC method, and coherence- and mutual information-based metrics fall in between. Results largely remain similar when GSR and bandpass filtering are not applied, with the caveat that all FC methods other than partial correlation perform equally badly without the application of GSR (Figure S1). If only the absolute values of edge weights are taken or if negative edges are set to zero, the performance of Pearson correlation improves, but is still worse than other measures (Figure S2). Figure 3C shows the distribution of QC-FC correlations for each FC estimation method. The distribution for Pearson correlation is wider than for other methods, confirming that more edges are significantly associated with motion.

**Figure 3.**
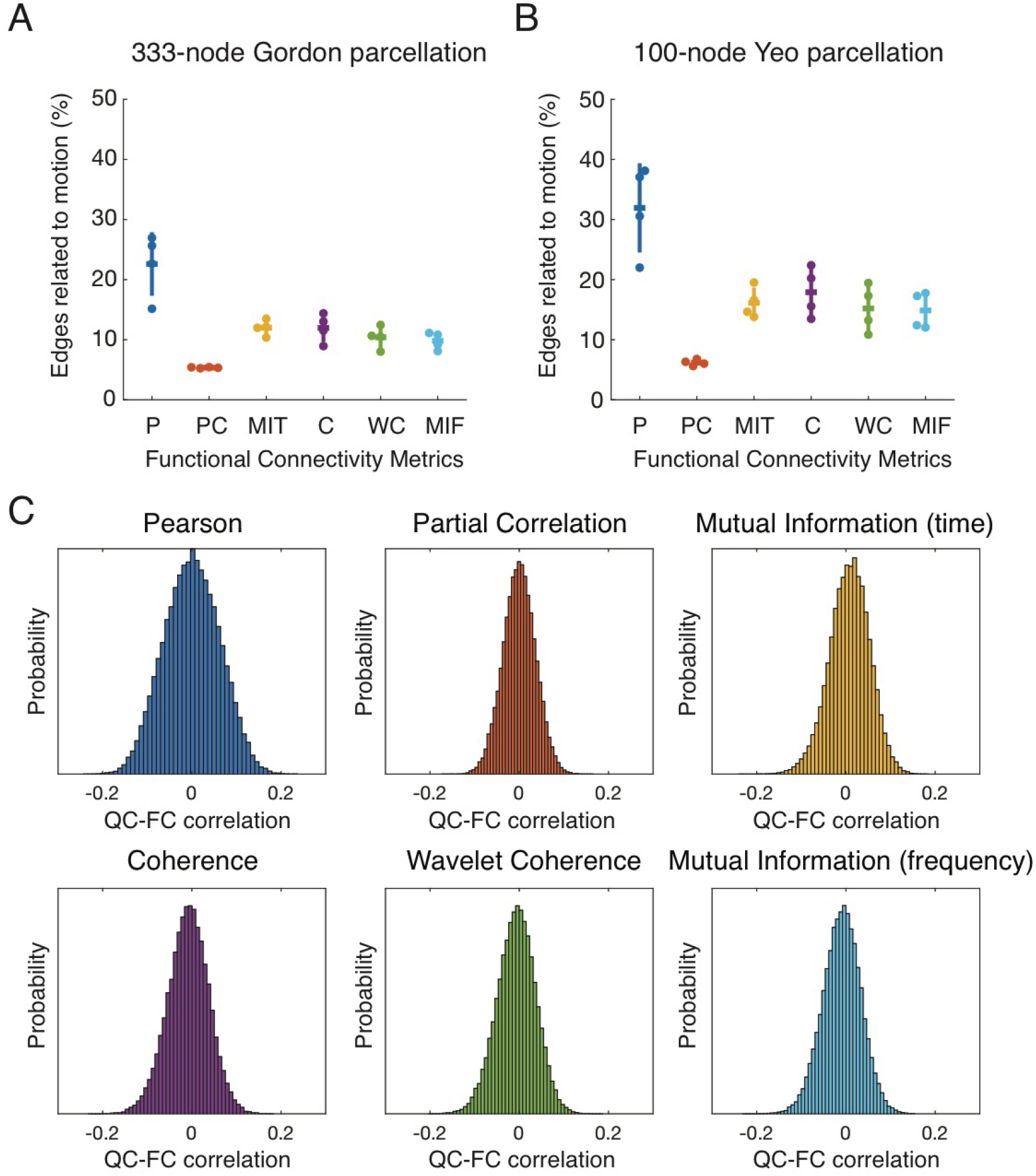
Pearson correlation shows higher residual QC-FC correlations. Fraction of edges significantly associated with motion for all 4 resting-state scans. **(A)** FC estimated using the 333-node Gordon parcellation. **(B)** FC estimated using the 100-node Schaefer parcellation. In both panels **A** and **B**, metric names are shortened as follows: P=Pearson, PC=Partial Correlation, MIT=Mutual Information (time), C=Coherence, WC=Wavelet Coherence, and MIF=Mutual Information (frequency). Notches represent the mean and error bars show the standard deviation. **(C)** Histograms of QC-FC correlations for all estimation methods. Note the wider distributions for edge weights computed using Pearson correlation.

### Motion differentially affects putative cognitive systems

Next, we analyzed the amount of motion artifact in edges connecting regions within specific putative cognitive systems. Figure 4 shows heatmaps of QC-FC correlations for all edges in the Gordon parcellation, arranged by the associated *a priori* defined systems^38^. Heatmaps of QC-FC correlations averaged for each system pair are shown in Figure S3. We also computed pairwise inter- and intra-community QC-FC correlations and rank-ordered them by their median values. The six highest ranked inter-community QC-FC correlations are shown in Figure 5. QC-FC correlations only within intra-system connections are shown in Figure S4.

**Figure 4.**
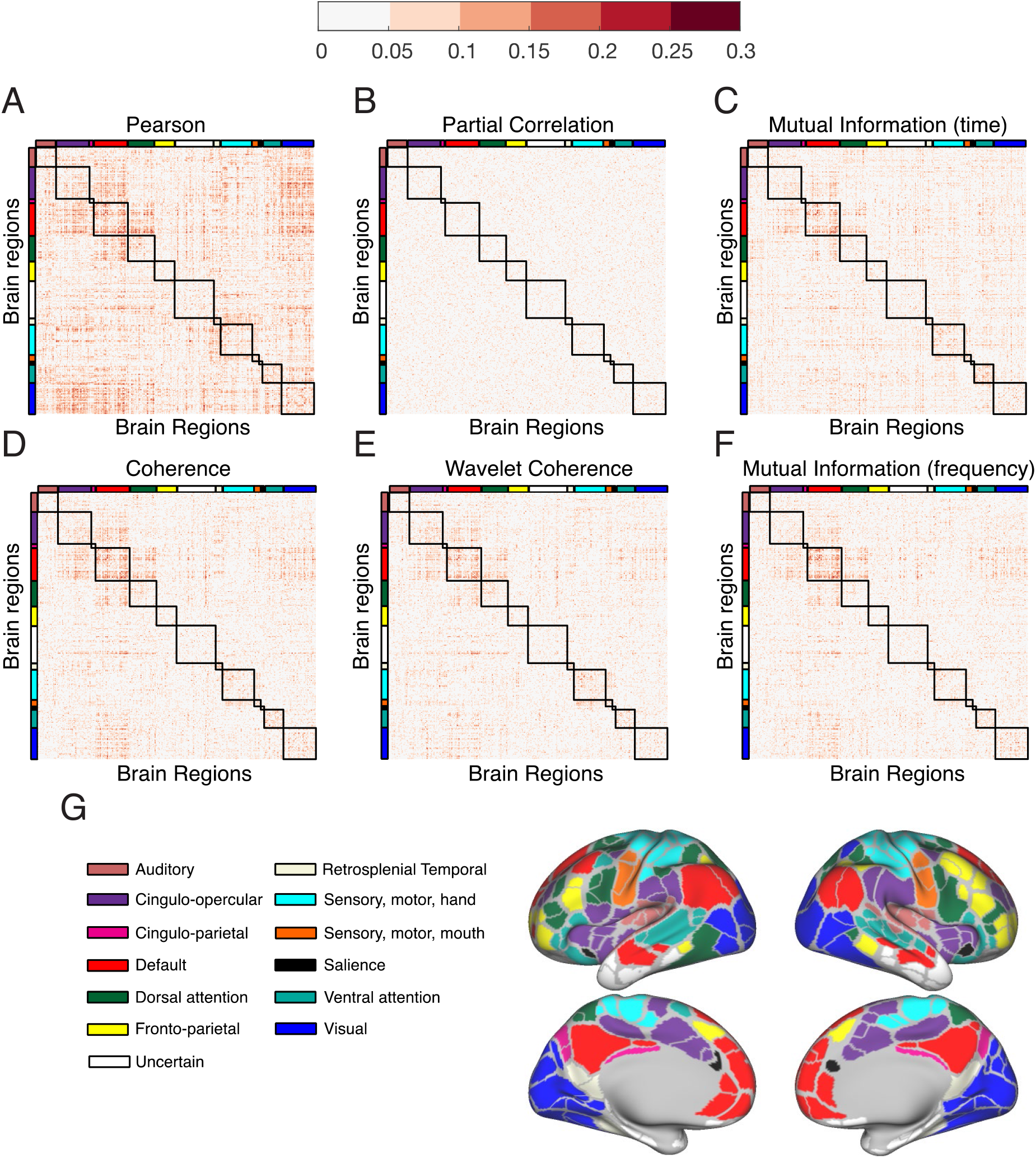
Distribution of QC-FC correlations across functional brain networks. QC-FC correlation heatmaps are shown for the REST1_LR scan, for the 333-node Gordon parcellation, and with canonical *a priori* defined systems labeled as colored bars. Each entry in the heatmap is the absolute value of the QC-FC correlation (across subjects), with edge weights estimated using **(A)** Pearson correlation, **(B)** partial correlation, **(C)** mutual information (time), **(D)** coherence, **(E)** wavelet coherence, and **(F)** mutual information (frequency). **(G)** Canonical systems for the Gordon parcellation displayed on the HCP S1200 group average cortical surface.

**Figure 5.**
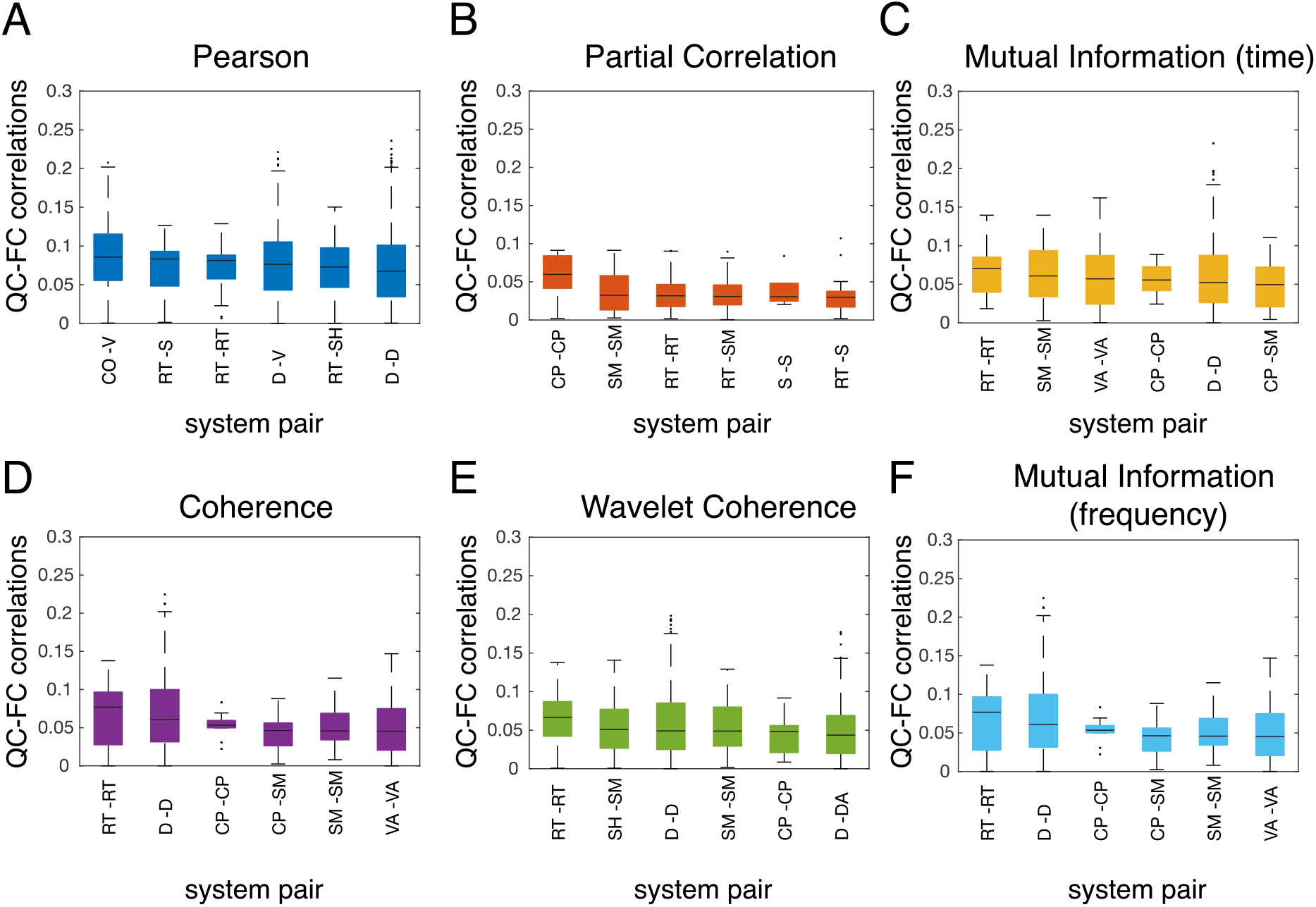
System affiliations of edges highly impacted by motion. Boxplots of inter-system QC-FC correlations are shown for the REST1_LR scan and the 333-node Gordon parcellation. All inter-system edges were rank-ordered by their median absolute QC-FC correlation, and the top 6 ranking distributions are shown for each FC metric. CO=cingulo-opercular; CP=cingulo-parietal; D=default; DA=dorsal attention; RT=retrosplenial temporal; SH=sensory, motor, hand; SM=sensory, motor, mouth; S=salience, VA=ventral attention; V=visual.

Our analysis reveals a number of interesting details about the differential vulnerability of brain systems to motion artifact. Edges within the default mode (D-D) and the retrosplenial temporal systems (RT-RT) appear to be especially vulnerable to motion artifact in most FC estimation methods except partial correlation (Figure 5, Figure S4). Notably, the fronto-parietal and auditory systems do not feature in the top six inter-system QC-FC correlations for any FC measure. For Pearson correlation, edges between cingulo-opercular and visual (CO-V), and between default and visual (D-V) systems have high QC-FC correlations but are not affected as much in the other FC measures.

### Distance-dependence of motion artifact

Next, we evaluated the distance-dependence of motion artifact by measuring edgewise correlations between the Euclidean distance between nodes and the edge’s absolute QC-FC correlation value. We find that edges estimated using Pearson correlation have higher positive distance-dependence than other methods, implying that long-distance edges are more affected by motion than short-distance edges (Figure 6). It is to be noted that the distance-dependence of QC-FC correlations is overall quite low^17^, which likely reflects the stringent preprocessing protocol employed. Results without application of GSR and bandpass filtering are shown in Figure S5.

**Figure 6.**
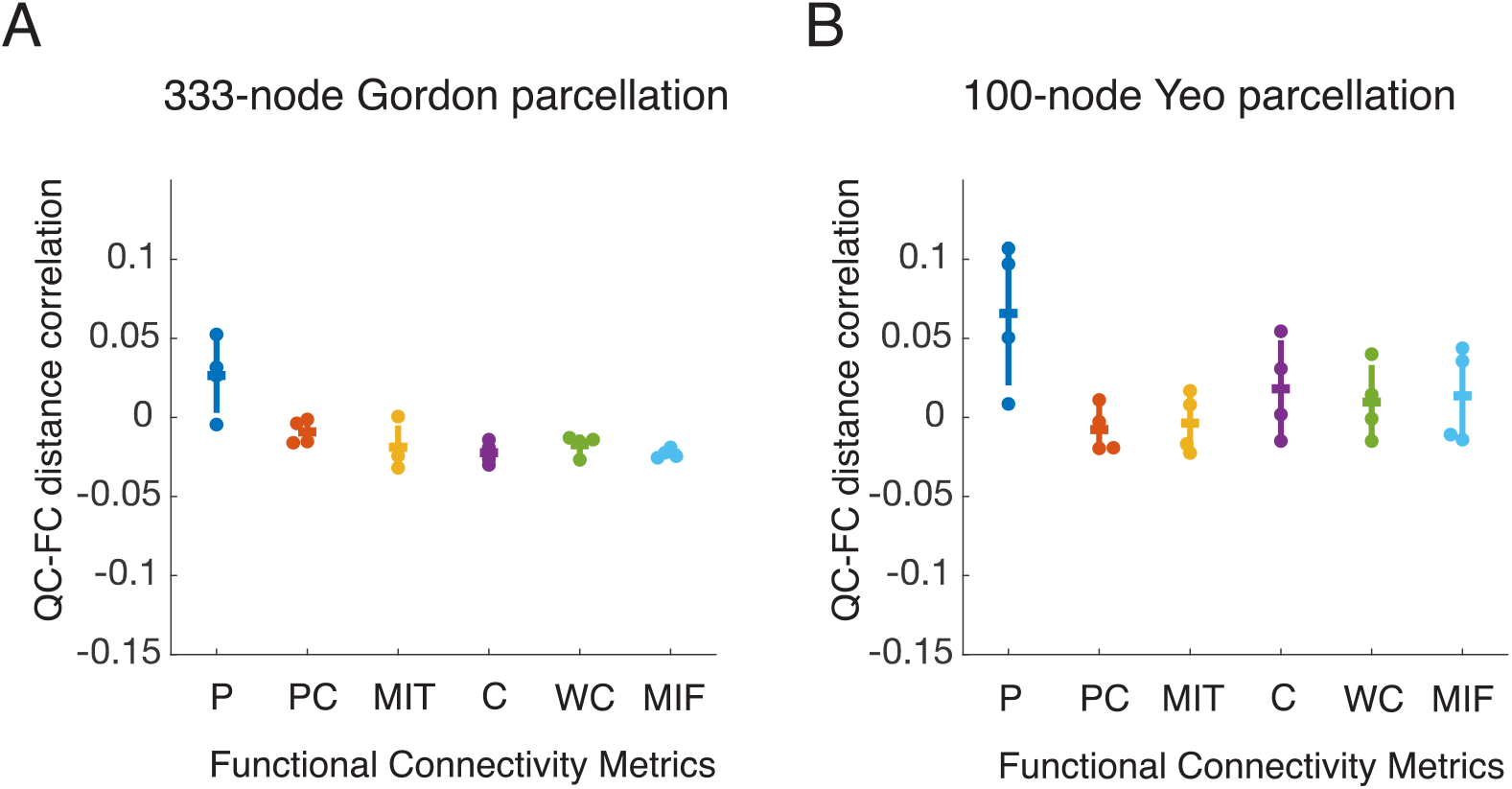
Distance-dependence of QC-FC correlations. Residual distance-dependence of motion artifact for different FC estimation methods, shown for all 4 resting-state scans. **(A)** FC estimated using the 333-node Gordon parcellation. **(B)** FC estimated using the 100-node Schaefer parcellation. In both panels **A** and **B**, the functional connectivity metric names are abbreviated as follows: P=Pearson, PC=Partial Correlation, MIT=Mutual Information (time), C=Coherence, WC=Wavelet Coherence, and MIF=Mutual Information (frequency). Notches represent the mean and error bars show the standard deviation.

### Test-retest reliability of functional connectivity

To estimate the reproducibility in functional connectivity estimates with different methods, we measured the intra-class correlation across 4 resting state scans in the HCP dataset. Panels A and B of Figure 7 show that the intra-class correlations are highest for edges estimated using Pearson correlation and lowest for edges estimated using partial correlation. The relatively high reliability of Pearson correlation edges could be due to accurate estimates of trait-like biology or could be due to a sensitivity to a highly reliable third-party variable, such as motion. To determine whether the latter could be the case, we evaluated the reliability of subject motion. We found that the intra-class correlation for relative RMS motion was also high (0.72), indicating that motion itself is reproducible across scans. In order to separate the reproducibility of motion from reproducibility of FC edges, we re-computed the intra-class correlations for edges that were in the bottom 20% of absolute QC-FC correlation values in all 4 scans. This analysis showed that results remained largely similar after mitigating the influence of reliable motion, indicating that edges estimated using Pearson correlation are more reproducible over this time scale than edges estimated using other methods (Figure 7C, D). Results remained largely similar without application of GSR and bandpass filtering (Figure S6).

**Figure 7.**
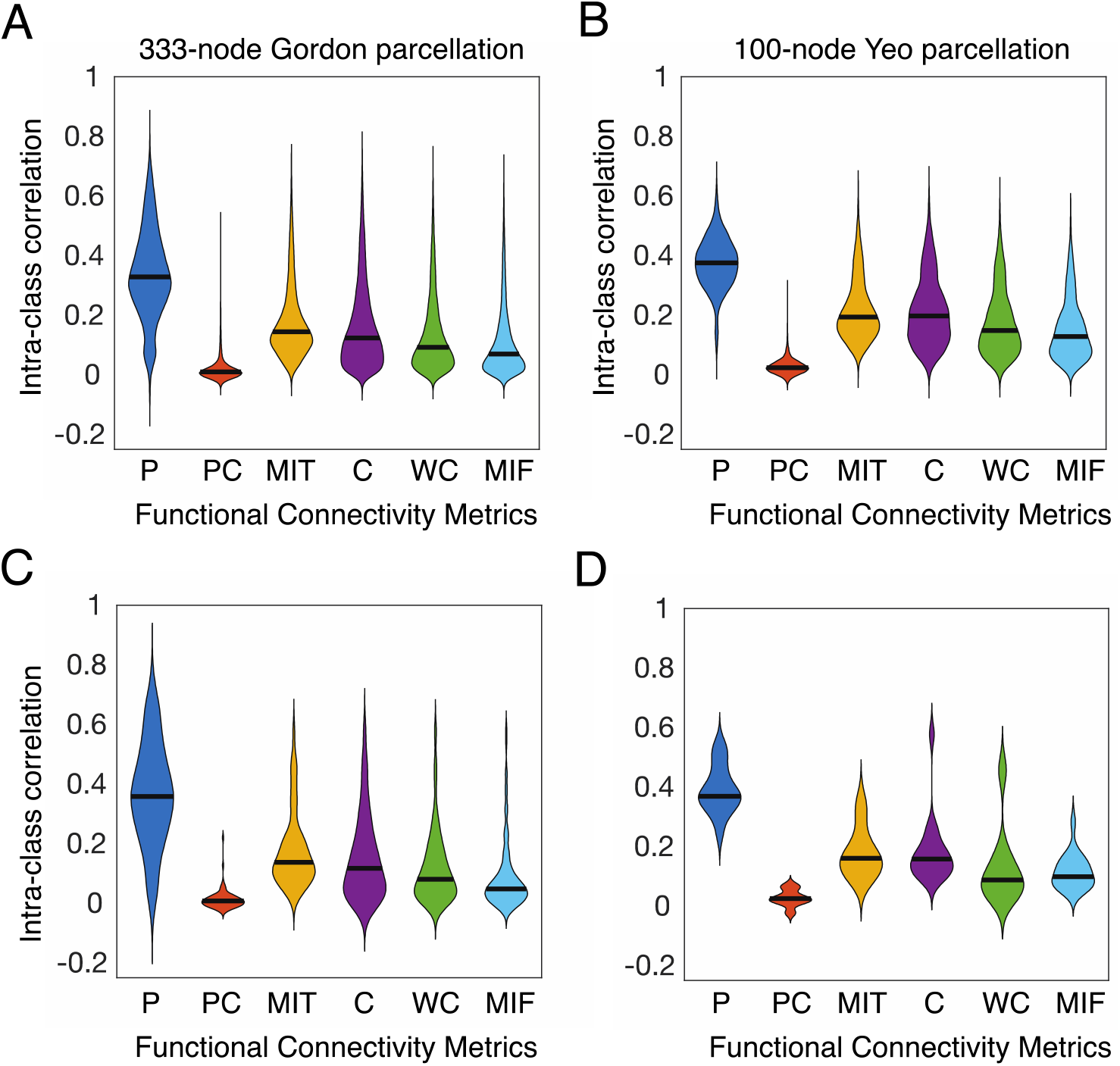
Test-retest reliability of edge weights for different FC metrics. (**A**-**B**) Intra-class correlation for all edge weights across 4 resting-state scans. **(A)** FC estimated using the 333-node Gordon parcellation. **(B)** FC estimated using the 100-node Schaefer parcellation. (**C-D**) Intra-class correlation for edges in the bottom 20% of QC-FC correlations for all 4 resting-state scans. **(C)** Gordon parcellation. **(D)** Schaefer parcellation. For all panels, the names of functional connectivity metrics have been abbreviated as follows: P=Pearson, PC=Partial Correlation, MIT=Mutual Information (time), C=Coherence, WC=Wavelet Coherence, and MIF=Mutual Information (frequency).

### System Identifiability

Finally, we examined the extent to which different metrics of FC resulted in different levels of system identifiability, or identification of the coherent intramodular structure that is commonly found in functional brain networks^48,49^. Panels A and B of Figure 8 show that on average, our measure of system idengifiability, the modularity quality index Q, is highest in systems estimated using Pearson’s correlation, lowest in all non-correlation based methods and intermediate for partial correlation.

**Figure 8.**
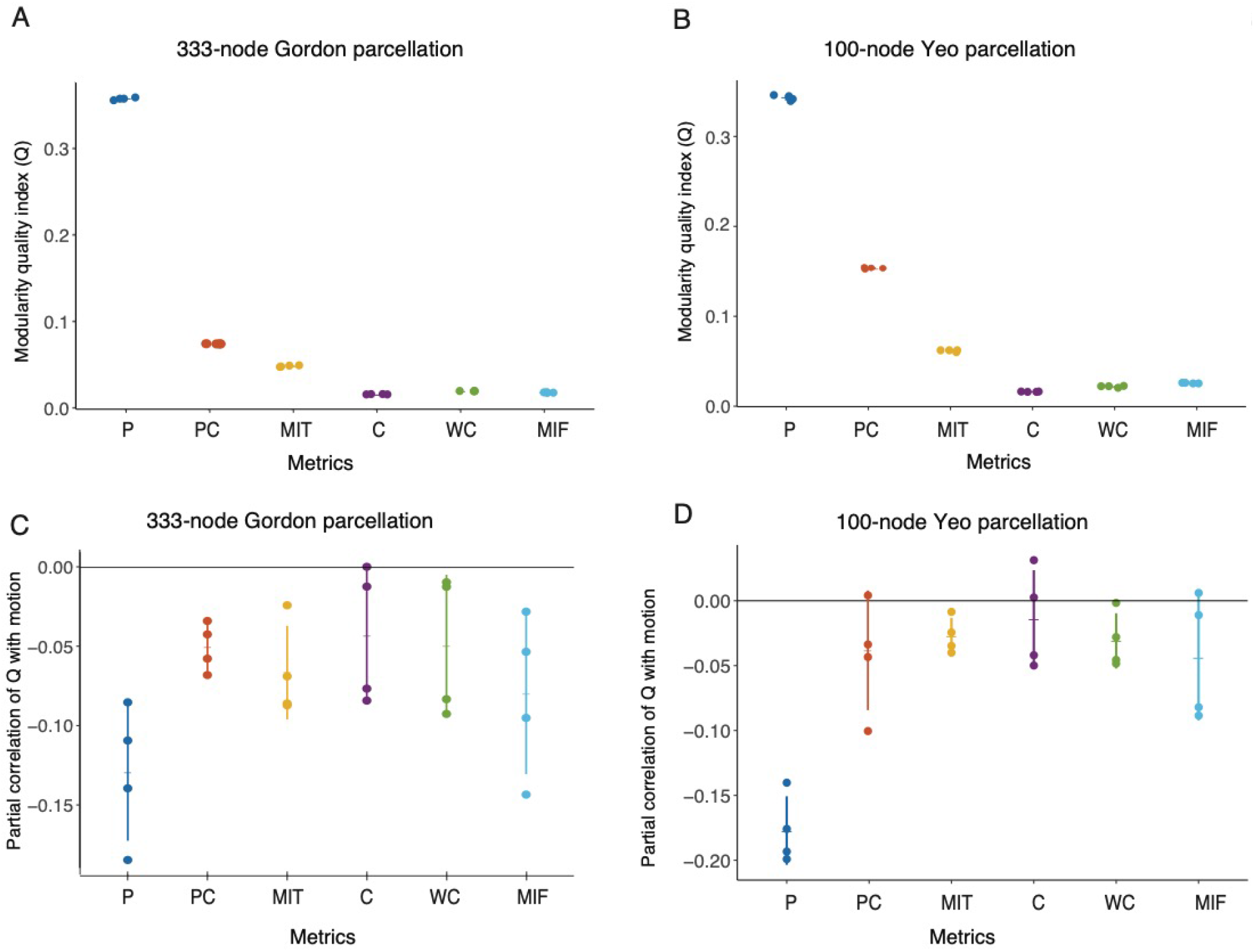
System identifiability is differentially associated with motion across FC metrics. **(A, B)** The modularity quality index *Q* is highest in networks estimated using Pearson correlation, lowest for non-correlation based metrics and intermediate for partial correlation. Modularity maximization was tuned across subjects to estimate between 6 and 8 communities for each metric. (**C, D)** Partial correlation between *Q* and mean relative RMS, controlling for average network weight, age, and sex. Edges estimated using Pearson correlation show large negative correlations between subject motion and *Q*. P=Pearson, PC=Partial Correlation, MIT=Mutual Information (time), C=Coherence, WC=Wavelet Coherence, MIF=Mutual Information (frequency). Notches represent the mean and error bars show the standard deviation.

The relatively high system identifiability in correlation-based metrics could be due to an accurate sensitivity to the biology of putative cognitive systems or could be due to motion impacting regions of the brain in a spatially heterogeneous manner that partially drives the data-driven identification of systems. To address these possibilities, we studied the relation between *Q* and motion. We find that the relationship between *Q* and motion is also strongest in Pearson correlation compared to all other FC metrics, even when controlling for average weight, age, and sex (Figure 8C, D). Results remained largely similar without application of bandpass filtering, but the relationship between *Q* and motion was similar among all FC methods without application of GSR (Figure S7, Figure S8).

We also estimated *Q* from networks containing only the absolute values of edge weights, which reduces but does not eliminate differences in system identifiability between correlation-based metrics and other metrics (Figure S9). Finally, to ensure that differences in system identifiability were not driven by differences in the functional systems detected when maximizing the modularity quality index, we also calculated *Q* using the canonical system partition associated with each of our 2 parcellations and obtained similar results (Figure S10).

To further ensure that results were not driven by variability in edge weight distributions across FC metrics, we examined two boundary cases from prior results as the FC metrics of interest: Pearson’s correlation and wavelet coherence. We reordered the edge weight values in our *weights* matrix (see Supplementary Methods) to reflect the rank order of weights in the ordering matrix. We estimated the modularity quality index Q of these reordered matrices, and found that Q was consistently higher when the ordering matrix was derived from Pearson’s correlations (Figure 9). Taken together, these results suggest that correlation-based FC metrics consistently result in higher levels of system identifiability, and that they may better reflect the modular architecture of functional brain networks than other methods.

**Figure 9.**
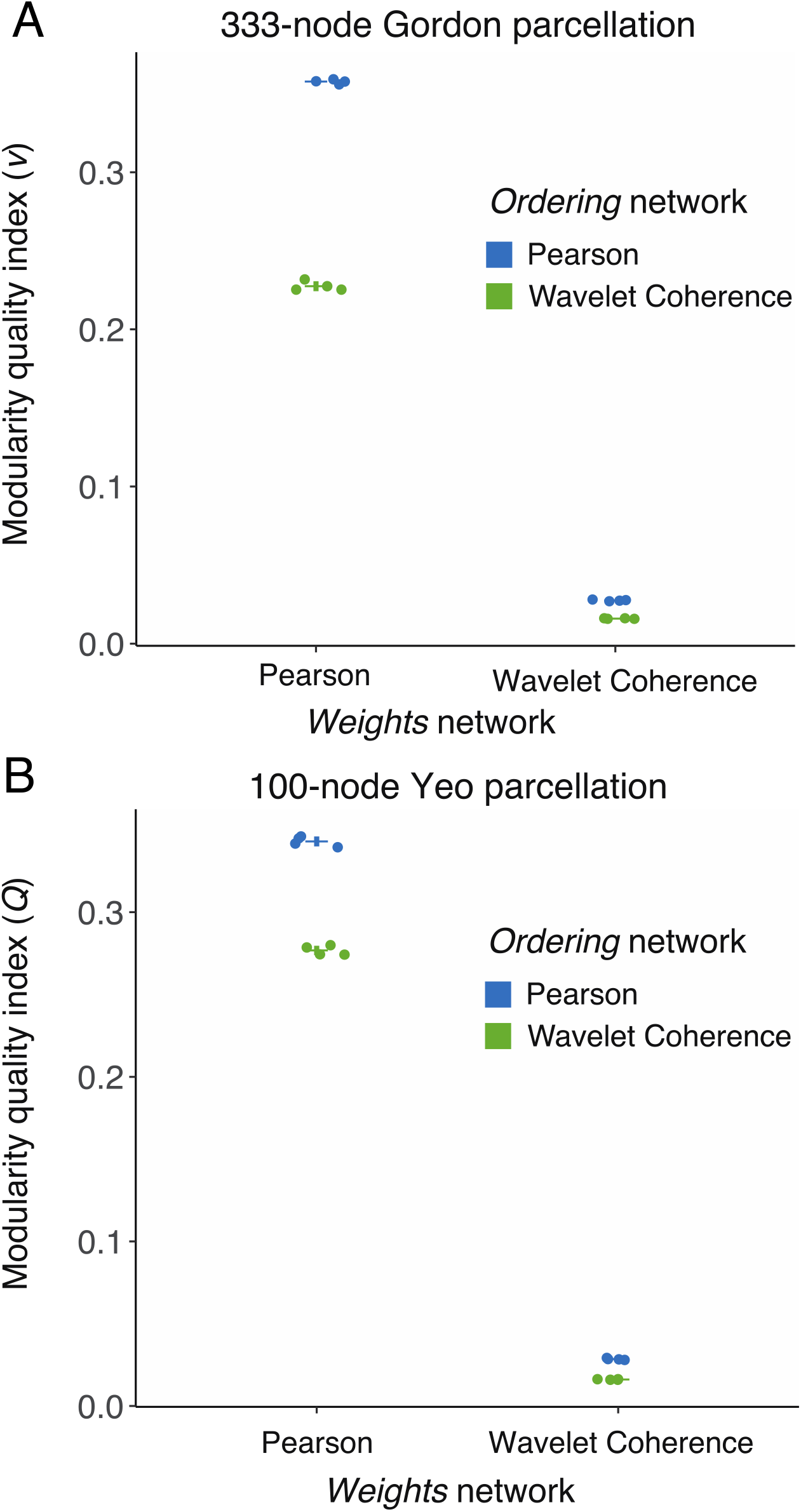
A correlation-based metric results in higher system identifiability even when using edge weight ranks rather than edge weight values. **(A, B)** Holding the edge weight distribution constant, higher *Q* is found when using the ranking of a time-based metric (Pearson) than a time-frequency-based metric (Wavelet Coherence). Notches represent the mean and error bars show the standard deviation.

## Discussion

In this report, we systematically investigated the sensitivity to motion of six different FC estimation measures drawn from the correlation, coherence, and mutual information families, based on their performance on commonly used benchmarks. The context, implications, and limitations of our results are discussed below.

### Clear distinction between correlation-based FC measures and other measures

Our main finding is that FC edges estimated using Pearson correlation result in a high fraction of edges significantly correlated with motion and a relatively high distance-dependence of motion artifact compared to all other methods. By contrast, using partial correlation almost completely eliminated the relationship between estimated edge weights and motion as well as their distance-dependence. These results were largely maintained when global signal regression and bandpass filtering were not applied.

Head motion artifact predominantly manifests as spurious signal fluctuations in BOLD signal across multiple voxels in the brain^16,30^. Since traditional functional connectivity based on full correlation measures temporal covariance, it follows that such measures of connectivity are directly impacted by artifactual covariance introduced by head motion. Partial correlation estimates, on the other hand, measure the temporal covariance between time series after regressing out all other time series in the network, thereby eliminating shared covariance induced by head motion artifact. Among the other FC methods we evaluated, coherence-based FC measures quantify statistical dependencies in the frequency domain, including phase locking and correlation in power spectra. We also evaluated mutual information in the frequency domain^27^, which is an information theoretic measure of relationships in the frequency domain. Frequency-based FC measures are less likely to be influenced by short-lived temporal fluctuations in the BOLD signal. Taken together, our findings indicate that the statistical properties of full correlation render them relatively more sensitive to temporal outliers introduced by head motion, an effect that is reduced by using frequency-based connectivity estimation, and effectively eliminated by regressing out common sources of artifact through partial correlation.

In our study, we averaged frequency-based connectivity estimates (coherence, wavelet coherence, and mutual information in frequency) within a low frequency band (0.009-0.08Hz). Although information on the power spectral properties of motion artifact is limited, some prior studies have shown that motion affects the spectral power and connectivity estimates mainly at high frequencies^55–57^. It is therefore possible that averaging connectivity estimates within a low frequency band reduced the impact of high-frequency motion artifact in these measures. Further, if motion artifact manifested in any one given frequency, the process of averaging in multiple frequency bands may have diluted the overall impact of motion on the FC estimates.

We found that the performance of Pearson correlation on the QC-FC benchmark improved when taking the absolute values of edge weights, and when negative edges were set to zero (Figure S2). Analysis of fully connected complex networks with positive and negative weights can be rigorously performed^50^. However, the interpretation of negative correlations is controversial, especially in the context of global signal regression^58–60^. As a result, many studies omit negative edges from analyses of functional and dynamic connectivity^61,62^. Our results indicate that omitting negative edges or taking their absolute values might also reduce the susceptibility to motion artifact, although it is also possible that this step decrements sensitivity to individual differences.

The systems whose edges were most affected by motion differed among FC estimation methods. For instance, connections between large-scale systems such as the default mode, cingulo-opercular and visual systems were related to motion in Pearson correlation but not in other FC methods. In contrast, within-system edges in the default mode system and retrosplenial temporal cortex were strongly related to motion in all FC estimation methods other than partial correlation. This last result deserves attention, given the large number of scientific hypotheses surrounding the brain’s default mode system^54,63^.

The strong relationship between motion and default mode connectivity is unlikely solely due to geometry because other networks whose edges are less correlated with motion, including the frontoparietal system, are similarly distributed with anterior and posterior nodes on both medial and lateral surfaces. It is possible that individuals with specific patterns of default mode connectivity find it more difficult to remember to stay still as their minds wander^64–66^.

Retrosplenial cortex, another system that we found to have a strong relationship with motion, is often functionally integrated with the default mode system to support memory processes, but also plays a role in spatial navigation and locomotion^67–69^. Specifically, retrosplenial cortex integrates vestibular input, which encodes head position, with visual cortex, to calibrate self-motion with visual motion signals^70,71^. Our findings highlight the need to carefully consider the confounding effects of motion, as well as the causes of motion, while studying these systems.

Significantly, Pearson correlation scored highest and partial correlation lowest on test-retest reliability. While the high reliability of edges estimated using Pearson correlation was partly caused by the well-known reproducibility of motion itself^72,73^, we observed similar results when restricting our analysis to the 20% of edges with the lowest absolute QC-FC correlations for all 4 scans. This additional finding suggests that Pearson correlation, while highly sensitive to motion, leads to relatively reproducible functional connectivity estimates. On the other hand, edges estimated using partial correlation, while not associated with motion, vary highly between scans.

Perhaps surprisingly, Pearson correlation resulted in significantly higher system identifiability than other measures. This observation suggests that findings of higher system identifiability in full correlation methods are not solely driven by motion, and that these metrics, while highly motion-sensitive, might excel in detecting coherent community structure. Alternatively, these findings may hint that the well-established finding of a modular architecture in human functional brain networks may be relatively metric-dependent^74,75^.

The success with which correlation-based measures detect modular architecture may be due to the presence of negative edge weights or anticorrelations in these measures, which contribute to reducing inter-module connections in calculations of modularity quality. A negative edge calculated using Pearson correlation, for instance between the default mode and dorsal attention systems, reduces the overall inter-module connectivity, crystallizing the boundaries between modules. However, the same edge calculated using coherence is highly positive, increasing inter-module connectivity and obfuscating boundaries between modules. Further, taking absolute values of correlation-based edges acts as a similar transformation, converting negative edges to positive edges, reducing the modularity quality index. The presence of negative edges or anticorrelation between internal and external attention systems has been argued to reflect a functional toggle between systems^59,76–79^. It is therefore unclear which method best captures true interactions between systems. Indeed, both could reflect complementary aspects of network dynamics if dorsal attention activation lags default mode activation at a consistent delay.

### Implications for researchers

We have shown that moving away from standard FC metrics based on full correlation can improve the robustness of FC estimates to head motion. However, Pearson correlation excels at detecting community structure and is highly reliable. Our findings indicate that the FC estimation method should be chosen carefully based on the nature of the study. For instance, studies on group comparisons, where motion artifact can introduce systematic bias in connectivity estimates^16^, could benefit from using frequency-based FC estimation methods like coherence. The appropriate choice for studies on modular brain architecture, in contrast, might remain Pearson correlation. Notably, partial correlation offers a best-of-both-worlds option – low contamination by motion artifact and relatively high system identifiability, with the caveat of low reliability. Our results also highlight a spatial heterogeneity in the impact of motion. FC edges in the default mode and retrosplenial cortex were especially sensitive to the effects of motion. Studies that explore the fine-scale organization and function of these networks could benefit from exploring different choices of FC estimation. Finally, Pearson correlation is computationally less expensive than the rest of the measures reported here and might therefore be preferable for large datasets where computational time is at a premium.

### Limitations

It is prudent to mention several limitations of our study. First, the lack of a noise-free ground truth is a challenge while estimating the impact of motion artifact in real fMRI data. It is difficult to separate out true signal from noise in fMRI data, a challenge further complicated by findings that head motion is a stable trait, and likely related to an individual’s physiology and neural dynamics^80,81^. Second, due to the lack of ground truth, it is necessary to rely on indirect benchmarks such as QC-FC correlations and QC-FC distance-dependence. Central to the computation of these benchmarks is the estimation of an average measure of head motion for the whole scan from realignment estimates^16^. This average measure can miss important spatiotemporal details of motion. Future studies could use voxel-wise displacement maps to extract more detailed information about motion and its impact on FC^56^. Third, some recent studies have highlighted the qualitative differences in motion estimates and physiological noise parameters from datasets with fast sampling rates compared to older datasets with larger TRs^82,83^. Thus, the estimates of head motion from realignment parameters, including those used in the current study, may need to be modified in the future for datasets with smaller TRs. Fourth, we used data from the Human Connectome Project that was preprocessed using ICA-FIX. This denoising approach has been shown to be particularly effective with HCP data^32^. In the future, it might be beneficial to investigate the effect of varying FC estimation methods with more noisy datasets with different denoising pipelines. Further, because we imposed a fairly stringent motion exclusion threshold, it is unclear whether our results generalize to samples with higher motion including pediatric, geriatric, or psychiatric samples. Finally, we did not evaluate FC estimation methods from many statistical families, including Bayes nets, Granger causality, and generalized synchronization. Future studies could investigate additional families of FC estimation methods omitted from this study.

## Citation diversity statement

Recent work in neuroscience and other fields has identified a bias in citation practices such that papers from women and other minorities are under-cited relative to the number of such papers in the field^84–89^. Here we sought to proactively consider choosing references that reflect the diversity of the field in thought, form of contribution, gender, and other factors. We obtained predicted gender of the first and last author of each reference by using databases that store the probability of a name being carried by a woman ^87,90^. By this measure (and excluding self-citations to the first and last authors of our current paper), our references contain 13.9% woman/woman, 10.1% man/woman, 22.8% woman/man, and 53.2% man/man. This method is limited in that a) names, pronouns, and social media profiles used to construct the databases may not, in every case, be indicative of gender identity and b) it cannot account for intersex, non-binary, or transgender people. We look forward to future work that could help us to better understand how to support equitable practices in science.

## Data/code availability statement

All data used in this manuscript comes from the open-source Human Connectome Project. Code associated with this manuscript is available at https://github.com/arunsm/motion-FC-metrics.git.

## Acknowledgements

We thank Dr. Linden Parkes for helpful discussions. U.A.T was supported by the National Science Foundation Graduate Research Fellowship. A.P.M. was supported by a Jacobs Foundation Early Career Research Fellowship and the National Institute on Drug Abuse (1R34DA050297-01). ASM was primarily supported by the Paul G. Allen Family Foundation and the Army Research Office (Falk W911NF-18-1-0244). DSB would also like to acknowledge the John D. and Catherine T. MacArthur Foundation, the ISI Foundation, the Army Research Laboratory (W911NF-10-2-0022), the Army Research Office (Bassett-W911NF-14-1-0679, Grafton-W911NF-16-1-0474), the National Science Foundation (BCS1631550, PHY-1554488, NCS-FO-1926829), the National Institute of Mental Health (2-R01-DC-009209-11, R01-MH112847, R01-MH107235, R21-M MH-106799, R01-MH-116920), and the National Institute of Child Health and Human Development (1R01HD086888-01). The content is solely the responsibility of the authors and does not necessarily represent the official views of any of the funding agencies.

## Author contributions

Arun S. Mahadevan: Methodology, Software, Formal Analysis, Visualization, Writing – Original Draft Preparation

Ursula Tooley: Methodology, Software, Formal Analysis, Visualization, Writing – Original Draft Preparation

Maxwell P. Bertolero: Data curation, Writing – Original Draft Preparation

Allyson P. Mackey: Supervision, Writing – Reviewing and Editing

Danielle S. Bassett: Conceptualization, Supervision, Formal Analysis, Writing – Reviewing and Editing

## References

1. Buckner, R. L., Krienen, F. M. & Yeo, B. T. T. Opportunities and limitations of intrinsic functional connectivity MRI. Nat. Neurosci. 16, 832–837 (2013).

2. Achard, S., Salvador, R., Whitcher, B., Suckling, J. & Bullmore, E. A resilient, low-frequency, small-world human brain functional network with highly connected association cortical hubs. J. Neurosci. 26, 63–72 (2006).

3. van den Heuvel, M. P. & Hulshoff Pol, H. E. Exploring the brain network: A review on resting-state fMRI functional connectivity. Eur. Neuropsychopharmacol. 20, 519–534 (2010).

4. Morgan, S. E., Achard, S., Termenon, M., Bullmore, E. T. & Vértes, P. E. Low-dimensional morphospace of topological motifs in human fMRI brain networks. Netw. Neurosci. 2, 285–302 (2017).

5. Vértes, P. E. et al. Simple models of human brain functional networks. Proc. Natl. Acad. Sci. U. S. A. 109, 5868–5873 (2012).

6. Achard, S. & Bullmore, E. Efficiency and cost of economical brain functional networks. PLoS Comput. Biol. 3, 0174–0183 (2007).

7. Gu, S. et al. Emergence of system roles in normative neurodevelopment. Proc. Natl. Acad. Sci. 112, 13681–13686 (2015).

8. Tooley, U. A. et al. Associations between Neighborhood SES and Functional Brain Network Development. Cereb. Cortex 30, 1–19 (2020).

9. Thomason, M. E. Development of Brain Networks In Utero: Relevance for Common Neural Disorders. Biological Psychiatry vol. 0 (2020).

10. Wheelock, M. D. et al. Sex differences in functional connectivity during fetal brain development. Dev. Cogn. Neurosci. 36, 100632 (2019).

11. van den Heuvel, M. I. et al. Hubs in the human fetal brain network. Dev. Cogn. Neurosci. 30, 108–115 (2018).

12. Morgan, S. E., White, S. R., Bullmore, E. T. & Vértes, P. E. A Network Neuroscience Approach to Typical and Atypical Brain Development. Biological Psychiatry: Cognitive Neuroscience and Neuroimaging vol. 3 754–766 (2018).

13. Vértes, P. E. & Bullmore, E. T. Annual research review: Growth connectomics - The organization and reorganization of brain networks during normal and abnormal development. J. Child Psychol. Psychiatry Allied Discip. 56, 299–320 (2015).

14. Finn, E. S. et al. Functional connectome fingerprinting: Identifying individuals using patterns of brain connectivity. Nat. Neurosci. 18, 1664–1671 (2015).

15. Gratton, C. et al. Functional Brain Networks Are Dominated by Stable Group and Individual Factors, Not Cognitive or Daily Variation. Neuron 98, 439–452.e5 (2018).

16. Power, J. D., Schlaggar, B. L. & Petersen, S. E. Recent progress and outstanding issues in motion correction in resting state fMRI. Neuroimage 105, 536–551 (2015).

17. Ciric, R. et al. Benchmarking of participant-level confound regression strategies for the control of motion artifact in studies of functional connectivity. Neuroimage 154, 174–187 (2017).

18. Parkes, L., Fulcher, B., Yücel, M. & Fornito, A. An evaluation of the efficacy, reliability, and sensitivity of motion correction strategies for resting-state functional MRI. Neuroimage 171, 415–436 (2018).

19. Caballero-Gaudes, C. & Reynolds, R. C. Methods for cleaning the BOLD fMRI signal. Neuroimage 154, 128–149 (2017).

20. Burgess, G. C. et al. Evaluation of Denoising Strategies to Address Motion-Correlated Artifacts in Resting-State Functional Magnetic Resonance Imaging Data from the Human Connectome Project. Brain Connect. 6, 669–680 (2016).

21. Friston, K. J. Functional and Effective Connectivity: A Review. Brain Connect. 1, 13–36 (2011).

22. Sun, F. T., Miller, L. M. & D’Esposito, M. Measuring interregional functional connectivity using coherence and partial coherence analyses of fMRI data. Neuroimage 21, 647–658 (2004).

23. Bullmore, E. et al. Wavelets and functional magnetic resonance imaging of the human brain. Neuroimage 23, 234–249 (2004).

24. Beauchene, C., Roy, S., Moran, R., Leonessa, A. & Abaid, N. Comparing brain connectivity metrics: a didactic tutorial with a toy model and experimental data. J. Neural Eng. 15, 56031 (2018).

25. Shannon, C. E. A Mathematical Theory of Communication. Bell Syst. Tech. J. 27, 379–423 (1948).

26. Quiroga, R. Q., Kraskov, A., Kreuz, T. & Grassberger, P. On the performance of different synchronization measures in real data: a case study on EEG signals. 65, 1–14 (2001).

27. Salvador, R., Suckling, J., Schwarzbauer, C. & Bullmore, E. Undirected graphs of frequency-dependent functional connectivity in whole brain networks. Philos. Trans. R. Soc. B Biol. Sci. 360, 937–946 (2005).

28. Smith, S. M. et al. Network modelling methods for FMRI. Neuroimage 54, 875–91 (2011).

29. Wang, H. E. et al. A systematic framework for functional connectivity measures. Front. Neurosci. 8, 1–22 (2014).

30. Ciric, R. et al. Mitigating head motion artifact in functional connectivity MRI. Nat. Protoc. 13, 2801–2826 (2018).

31. Van Essen, D. C. et al. The WU-Minn Human Connectome Project: An overview. Neuroimage 80, 62–79 (2013).

32. Salimi-Khorshidi, G. et al. Automatic denoising of functional MRI data: Combining independent component analysis and hierarchical fusion of classifiers. Neuroimage 90, 449–468 (2014).

33. Griffanti, L. et al. ICA-based artefact removal and accelerated fMRI acquisition for improved resting state network imaging. Neuroimage 95, 232–247 (2014).

34. Bertolero, M. A. et al. The network architecture of the human brain is modularly encoded in the genome. (2019).

35. Schaefer, A. et al. Local-Global Parcellation of the Human Cerebral Cortex from Intrinsic Functional Connectivity MRI. Cereb. Cortex 28, 3095–3114 (2018).

36. Whitaker, K. J. et al. Adolescence is associated with genomically patterned consolidation of the hubs of the human brain connectome. Proc. Natl. Acad. Sci. U. S. A. 113, 9105–9110 (2016).

37. Vértes, P. E. et al. Gene transcription profiles associated with inter-modular hubs and connection distance in human functional magnetic resonance imaging networks. Philos. Trans. R. Soc. B Biol. Sci. 371, (2016).

38. Gordon, E. M. et al. Generation and Evaluation of a Cortical Area Parcellation from Resting-State Correlations. Cereb. Cortex 26, 288–303 (2016).

39. Strehl, A. & Ghosh, J. Cluster Ensembles --- A Knowledge Reuse Framework for Combining Multiple Partitions. J. Mach. Learn. Res. 3, 583–617 (2002).

40. Zhou, D., Thompson, W. K. & Siegle, G. MATLAB toolbox for functional connectivity. Neuroimage 47, 1590–1607 (2009).

41. Zhang, Z., Telesford, Q. K., Giusti, C., Lim, K. O. & Bassett, D. S. Choosing wavelet methods, filters, and lengths for functional brain network construction. PLoS One 11, 1–24 (2016).

42. Grinsted, A., Moore, J. C. & Jevrejeva, S. Application of the cross wavelet transform and wavelet coherence to geophysical time series. Nonlinear Process. Geophys. 11, 561–566 (2004).

43. Bassett, D. S. & Sporns, O. Network neuroscience. Nat. Neurosci. 20, 353–364 (2017).

44. Satterthwaite, T. D. et al. Impact of in-scanner head motion on multiple measures of functional connectivity: Relevance for studies of neurodevelopment in youth. Neuroimage 60, 623–632 (2012).

45. Shrout, P. E. & Fleiss, J. L. Intraclass correlations: Uses in assessing rater reliability. Psychol. Bull. 86, 420–428 (1979).

46. Newman, M. Modularity and community structure in networks. Proc. Natl. Acad. … 103, 8577–82 (2006).

47. Girvan, M. & Newman, M. E. J. Community structure in social and biological networks. Proc. Natl. Acad. Sci. U. S. A. 99, 7821–7826 (2002).

48. Porter, M. A., Onnela, J.-P. & Mucha, P. J. Communities in Networks. Not. AMS 56, 1082–1097 (2009).

49. Fortunato, S. Community detection in graphs. Phys. Rep. 486, 75–174 (2010).

50. Rubinov, M. & Sporns, O. Weight-conserving characterization of complex functional brain networks. Neuroimage 56, 2068–2079 (2011).

51. Traag, V. A. & Bruggeman, J. Community detection in networks with positive and negative links. Phys. Rev. E - Stat. Nonlinear, Soft Matter Phys. 80, 1–6 (2009).

52. Gómez, S., Jensen, P. & Arenas, A. Analysis of community structure in networks of correlated data. Phys. Rev. E - Stat. Nonlinear, Soft Matter Phys. 80, 1–5 (2009).

53. Blondel, V. D., Guillaume, J. L., Lambiotte, R. & Lefebvre, E. Fast unfolding of communities in large networks. J. Stat. Mech. Theory Exp. 2008, 0–12 (2008).

54. Buckner, R. L. & DiNicola, L. M. The brain’s default network: updated anatomy, physiology and evolving insights. Nat. Rev. Neurosci. (2019) doi:10.1038/s41583-019-0212-7.

55. Salvador, R. et al. A simple view of the brain through a frequency-specific functional connectivity measure. Neuroimage 39, 279–289 (2008).

56. Satterthwaite, T. D. et al. An improved framework for confound regression and filtering for control of motion artifact in the preprocessing of resting-state functional connectivity data. Neuroimage 64, 240–256 (2013).

57. Kim, J., Van Dijk, K. R. A., Libby, A. & Napadow, V. Frequency-dependent relationship between resting-state functional magnetic resonance imaging signal power and head motion is localized within distributed association networks. Brain Connect. 4, 30–39 (2014).

58. Anderson, J. S. et al. Network anticorrelations, global regression, and phase-shifted soft tissue correction. Hum. Brain Mapp. 32, 919–934 (2011).

59. Chai, X. J., Castañán, A. N., Öngür, D. & Whitfield-Gabrieli, S. Anticorrelations in resting state networks without global signal regression. Neuroimage 59, 1420–1428 (2012).

60. Murphy, K. & Fox, M. D. Towards a consensus regarding global signal regression for resting state functional connectivity MRI. Neuroimage 154, 169–173 (2017).

61. Grady, C., Sarraf, S., Saverino, C. & Campbell, K. Age differences in the functional interactions among the default, frontoparietal control, and dorsal attention networks. Neurobiol. Aging 41, 159–172 (2016).

62. Chan, M. Y., Park, D. C., Savalia, N. K., Petersen, S. E. & Wig, G. S. Decreased segregation of brain systems across the healthy adult lifespan. Proc. Natl. Acad. Sci. U. S. A. 111, E4997–E5006 (2014).

63. Fox, M. D. & Raichle, M. E. Spontaneous fluctuations in brain activity observed with functional magnetic resonance imaging. Nat. Rev. Neurosci. 8, 700–11 (2007).

64. Christoff, K., Gordon, A. M., Smallwood, J., Smith, R. & Schooler, J. W. Experience sampling during fMRI reveals default network and executive system contributions to mind wandering. Proc. Natl. Acad. Sci. U. S. A. 106, 8719–8724 (2009).

65. Kajimura, S., Kochiyama, T., Nakai, R., Abe, N. & Nomura, M. Causal relationship between effective connectivity within the default mode network and mind-wandering regulation and facilitation. Neuroimage 133, 21–30 (2016).

66. Golchert, J. et al. Individual variation in intentionality in the mind-wandering state is reflected in the integration of the default-mode, fronto-parietal, and limbic networks. Neuroimage 146, 226–235 (2017).

67. Vann, S. D., Aggleton, J. P. & Maguire, E. A. What does the retrosplenial cortex do? Nat. Rev. Neurosci. 10, 792–802 (2009).

68. Fischer, L. F., Mojica Soto-Albors, R., Buck, F. & Harnett, M. T. Representation of visual landmarks in retrosplenial cortex. Elife 9, 1–25 (2020).

69. Mao, D., Molina, L. A., Bonin, V. & McNaughton, B. L. Vision and Locomotion Combine to Drive Path Integration Sequences in Mouse Retrosplenial Cortex. Curr. Biol. 1–9 (2020) doi:10.1016/j.cub.2020.02.070.

70. Vélez-Fort, M. et al. A Circuit for Integration of Head- and Visual-Motion Signals in Layer 6 of Mouse Primary Visual Cortex. Neuron 98, 179–191.e6 (2018).

71. Chaplin, T. A. & Margrie, T. W. Cortical circuits for integration of self-motion and visual-motion signals. Curr. Opin. Neurobiol. 60, 122–128 (2020).

72. Noble, S., Scheinost, D. & Constable, R. T. A decade of test-retest reliability of functional connectivity: A systematic review and meta-analysis. Neuroimage 203, 116157 (2019).

73. van Dijk, K. R. A., Sabuncu, M. R. & Buckner, R. L. The influence of head motion on intrinsic functional connectivity MRI. Neuroimage 59, 431–438 (2012).

74. Bertolero, M. A., Thomas Yeo, B. T. & D’Esposito, M. The modular and integrative functional architecture of the human brain. Proc. Natl. Acad. Sci. U. S. A. 112, E6798–E6807 (2015).

75. Sporns, O. & Betzel, R. F. Modular Brain Networks. Annu. Rev. Psychol. 67, 613–640 (2016).

76. Gao, W. & Lin, W. Frontal parietal control network regulates the anti-correlated default and dorsal attention networks. Hum. Brain Mapp. 33, 192–202 (2012).

77. Clare Kelly, A. M., Uddin, L. Q., Biswal, B. B., Castellanos, F. X. & Milham, M. P. Competition between functional brain networks mediates behavioral variability. Neuroimage 39, 527–537 (2008).

78. Owens, M. M., Duda, B., Sweet, L. H. & MacKillop, J. Distinct functional and structural neural underpinnings of working memory. Neuroimage 174, 463–471 (2018).

79. Murphy, A. C., Bertolero, M. A., Papadopoulos, L., Lydon-Staley, D. M. & Bassett, D. S. Multiscale and multimodal network dynamics underpinning working memory. 3–6 (2019).

80. Engelhardt, L. E. et al. Children’s head motion during fMRI tasks is heritable and stable over time. Dev. Cogn. Neurosci. 25, 58–68 (2017).

81. Beyer, F. et al. Weight loss reduces head motion: Revisiting a major confound in neuroimaging. Hum. Brain Mapp. 1–5 (2020) doi:10.1002/hbm.24959.

82. Agrawal, U., Brown, E. N. & Lewis, L. D. Model-based physiological noise removal in fast fMRI. Neuroimage 116231 (2019) doi:10.1016/j.neuroimage.2019.116231.

83. Power, J. D. et al. Distinctions among real and apparent respiratory motions in human fMRI data. Neuroimage 201, 116041 (2019).

84. Caplar, N., Tacchella, S. & Birrer, S. Quantitative evaluation of gender bias in astronomical publications from citation counts. Nat. Astron. 1, (2017).

85. Chakravartty, P., Kuo, R., Grubbs, V. & McIlwain, C. #CommunicationSoWhite. J. Commun. 68, 254–266 (2018).

86. Dion, M. L., Sumner, J. L. & Mitchell, S. M. L. Gendered Citation Patterns across Political Science and Social Science Methodology Fields. Polit. Anal. 26, 312–327 (2018).

87. Dworkin, J. D. et al. The extent and drivers of gender imbalance in neuroscience reference lists. (2020).

88. Maliniak, D., Powers, R. & Walter, B. F. The gender citation gap in international relations. International Organization vol. 67 (2013).

89. Thiem, Y., Sealey, K. F., Ferrer, A. E., Trott, A. M. & Kennison, R. Just Ideas? The Status and Future of Publication Ethics in Philosophy: A White Paper. (2018).

90. Zhou, D. et al. Gender Diversity Statement and Code Notebook v1.0. (2020) doi:10.5281/zenodo.3672110.

