## Supplementary Information for "Evaluating the sensitivity of functional connectivity measures to motion artifact in resting-state fMRI data"

### Supplementary Methods

*System Identifiability*

To estimate the modularity quality index $Q$ for a given subject's functional brain network, we use a heuristic[1] to maximize $Q$ for a partition $M$ of nodes into $k$ communities. However, maximizing $Q$ while keeping the structural resolution parameter at $\gamma = 1$ may result in a different number of communities $k$ detected for different input data, across both subjects and FC metrics. Previous work examining the modularity quality index, $Q$, as a measure of system identifiability has failed to account for differences in the number of communities $k$ in the partition $M$.[2] In this work, we address this confound by tuning the structural resolution parameter $\gamma$ across subjects to estimate a partition $M$ that results in $6 < k < 8$ communities for each FC metric, based on prior evidence that cortical functional brain networks can be reliably and meaningfully divided into approximately seven communities[3]. In this fashion, we obtain a $\gamma$ for each metric that results in a similarly sized partition of nodes into $k$ communities.

Additionally, the average edge weight of a network can be highly associated with $Q$[4–6]. Thus, differences in the mean or distribution of edge weights obtained using different FC metrics (see Figure 1 in main text) might lead to interpretations of higher system identifiability, when in fact the difference in $Q$ is solely a product of differences in the distribution of edge weights. Previous work examining the modularity quality index, $Q$, as a measure of system identifiability has failed to account for the influence of edge weight distributions on $Q$.[2] In this work, we address this confound in two ways. First, in all analyses examining the associations between motion and $Q$, we control for the average edge weight of the network. Including the average weight as a covariate ensures that subsequent results reflect differences in network topology, rather than differences in average connectivity strength across FC metrics. We computed the partial correlation between the modularity quality index $Q$ and the relative mean RMS motion for each subject and for each FC metric, while controlling for average network weight, age, and sex. The resultant estimate served as a measure of the extent to which motion was associated with estimates of system identifiability, when accounting for differences in the average weight of the network, which varies across FC metrics.

Our first approach accounts only for variation in the average edge weight of the network, however, and not any variation in the *distribution* of edge weights across FC metrics. To further ensure that results are not driven by variability in edge weights across FC metrics, we developed an approach that preserves the topology of the network while ensuring that both the average edge weight and distribution of edge weights are the same. We used this approach to examine two boundary cases from prior results as the FC metrics of interest: Pearson correlation and wavelet coherence.

In this procedure, we designate a *weights* network, from which we extract actual edge weights, and an *ordering* network, from which we calculate the rank order of edge weights, from weakest to strongest. In this example, the network estimated from Pearson correlations will be our *weights* network, with the network resulting from wavelet coherence being our *ordering* network. We reorder the edge weight values in the matrix representing our *weights* network (Pearson) to match the rank ordering of edge weights in our *ordering* network (wavelet coherence). This process preserves the topology of the network contained in the *ordering* matrix while ensuring that both the average edge weight and distribution of edge weights are the same as those in the *weights* network. We then estimated $Q$, as above, using the $\gamma$ appropriate for the distribution of edge weight values (tuned for $6 < k < 8$ for the *weights* network). We compared this $Q$, estimated from the reordered network, to that of the original *weights* matrix as a measure of network identifiability.

Finally, to ensure that differences in system identifiability were not driven by differences in functional systems detected using modularity maximization, we also estimated $Q$ using the canonical partition for each parcellation. In these analyses, we did not employ a heuristic to maximize $Q$; rather, we used the *a priori* partition of parcels into systems that is associated with the Gordon 333-node and Schaefer 100-node parcellation[7,8]. For the Schaefer 100-node parcellation, we used the 17-system partition.

### Supplementary Figures

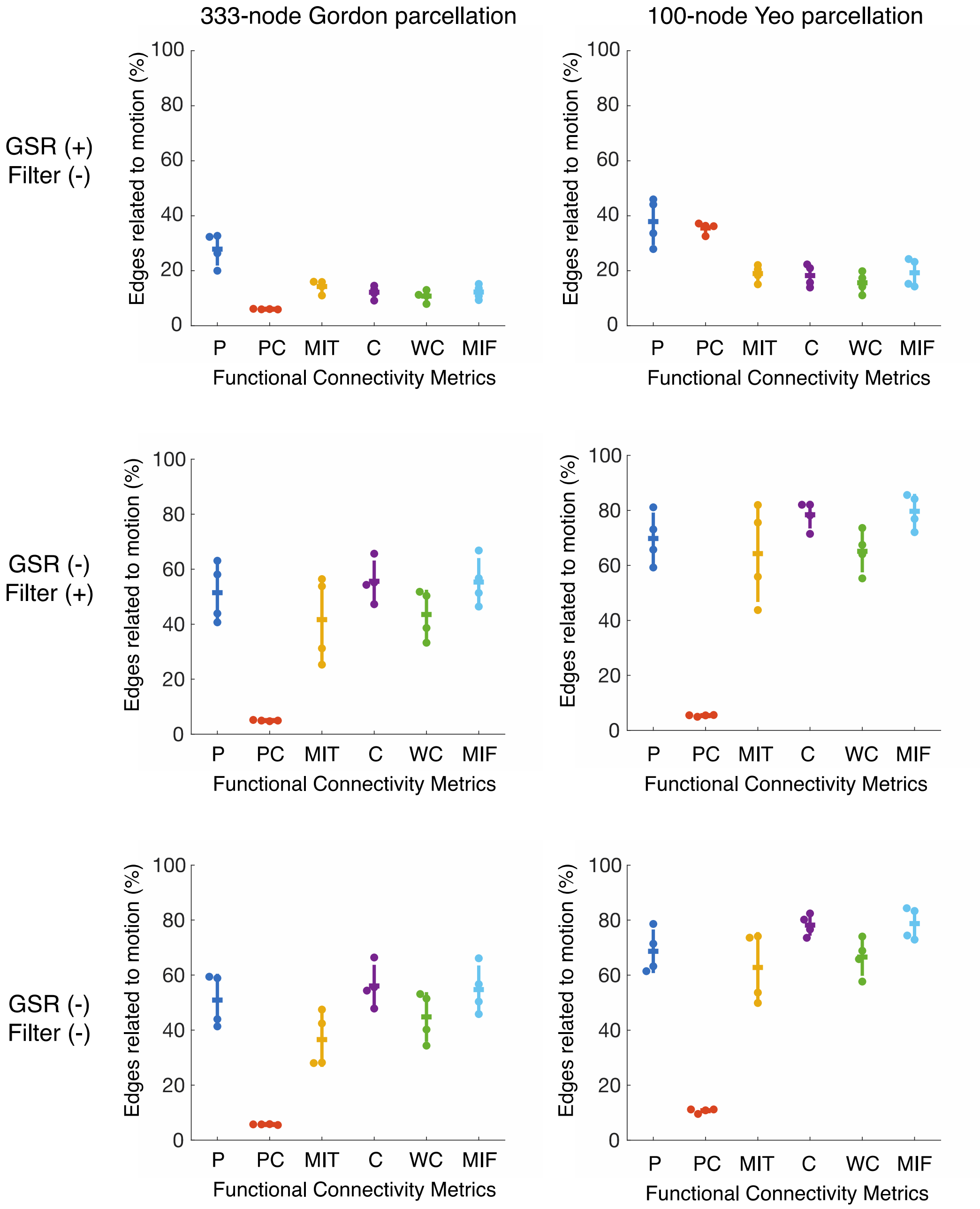


**Figure S1.** Fraction of edges significantly associated with motion for all 4 resting-state scans, with three different preprocessing variants defined by the inclusion or exclusion of global signal regression (GSR) and bandpass filtering. Results for 333-node parcellation provided by Gordon *et al.* (2016) shown in left column, and 100-node parcellation provided by Schaefer *et al.* (2017) shown in right column. P=Pearson, PC=Partial Correlation, MIT=Mutual Information (time), C=Coherence, WC=Wavelet Coherence, MIF=Mutual Information (frequency). Notches represent the mean of four resting-state scans and error bars show the standard deviation.

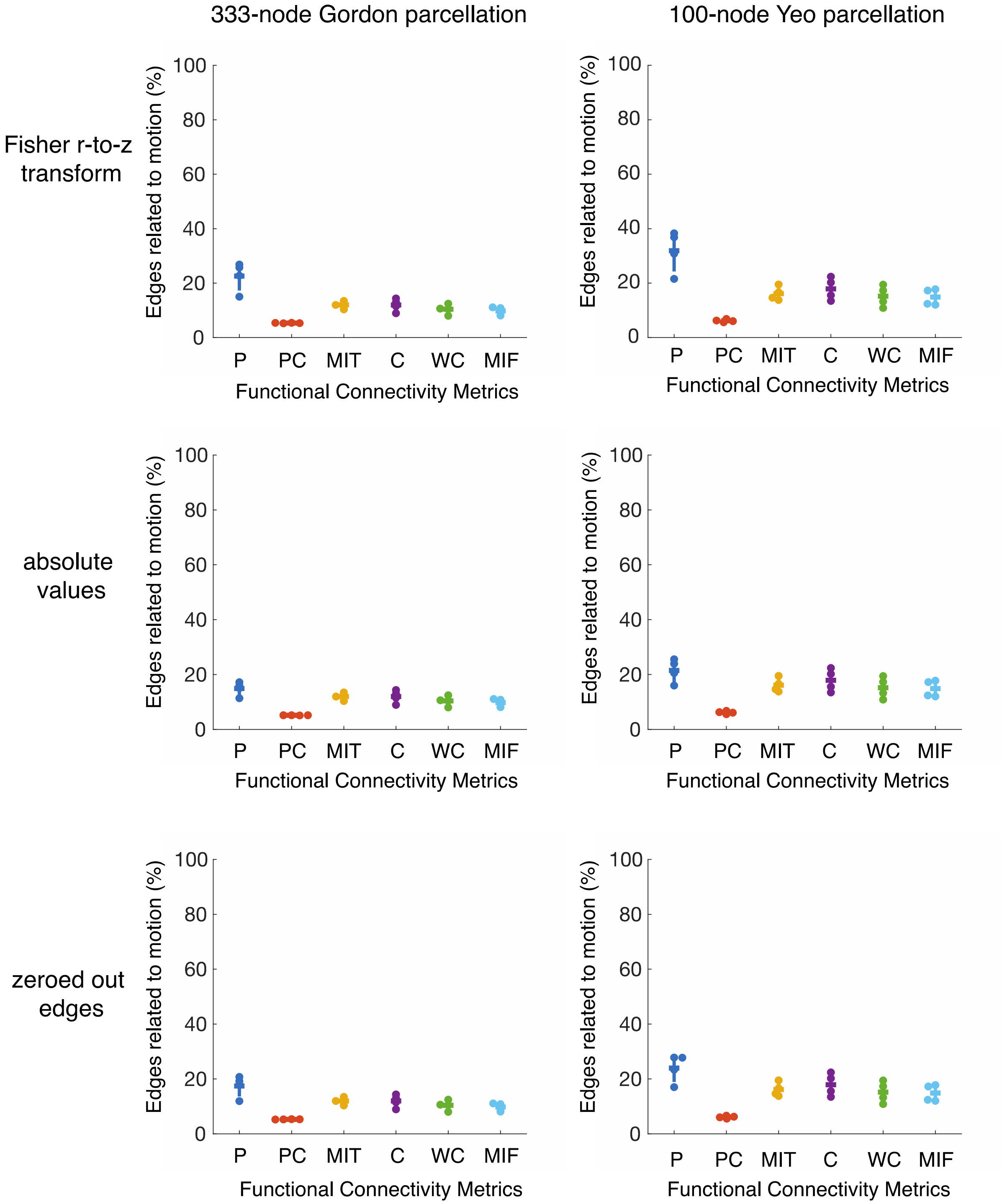


**Figure S2.** Fraction of edges significantly associated with motion for all 4 resting-state scans, with three different treatment of edges in correlation-based measures: application of Fisher r-to-z transform, taking absolute values of edges, and zeroing out of negative edges. Results for 333-node parcellation provided by Gordon *et al.* (2016) shown in left column, and 100-node parcellation provided by Schaefer *et al.* (2017) shown in right column. P=Pearson, PC=Partial Correlation, MIT=Mutual Information (time), C=Coherence, WC=Wavelet Coherence, MIF=Mutual Information (frequency).

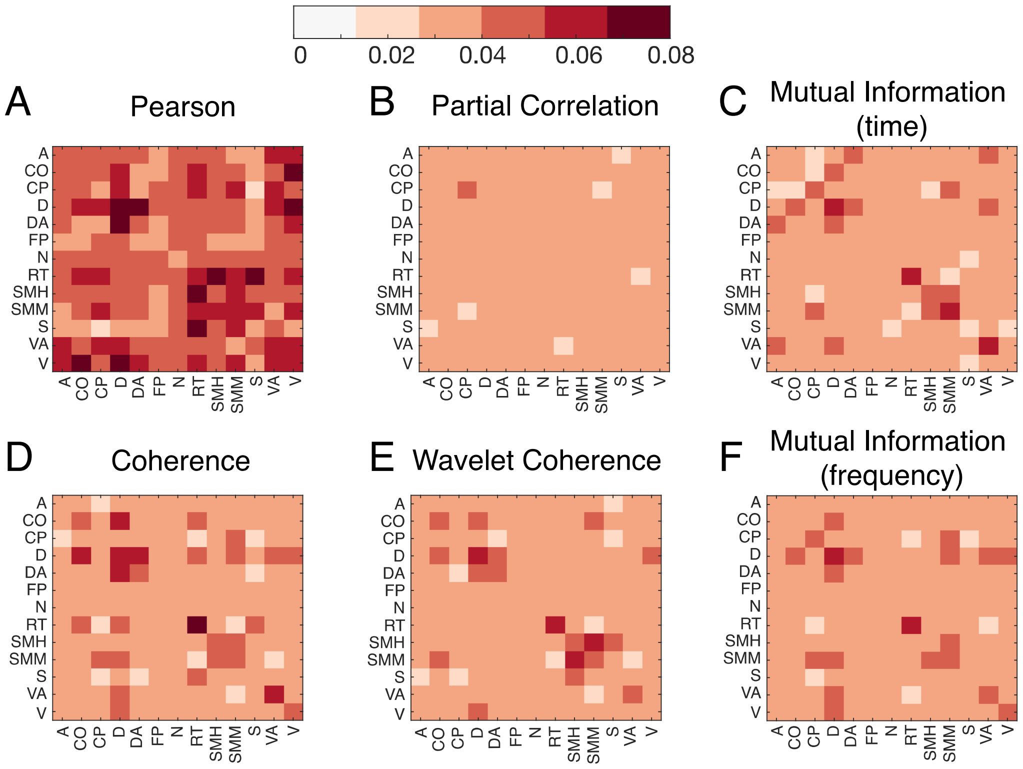


**Figure S3.** Average QC-FC correlation heatmaps are shown for the REST1_LR scan and the 333-node Gordon parcellation. Each entry in the heatmap is the average QC-FC correlation between the corresponding systems, with edge weights estimated using **(A)** Pearson correlation, **(B)** partial correlation, **(C)** mutual information (time), **(D)** coherence, **(E)** wavelet coherence, and **(F)** mutual information (frequency). A=auditory; CO=cingulo-opercular; CP=cingulo-parietal; D=default; DA=dorsal attention; FP=fronto-parietal; N=none; RT=retrosplenial temporal; SH=sensory, motor, hand; SM=sensory, motor, mouth; S=salience, VA=ventral attention; V=visual.

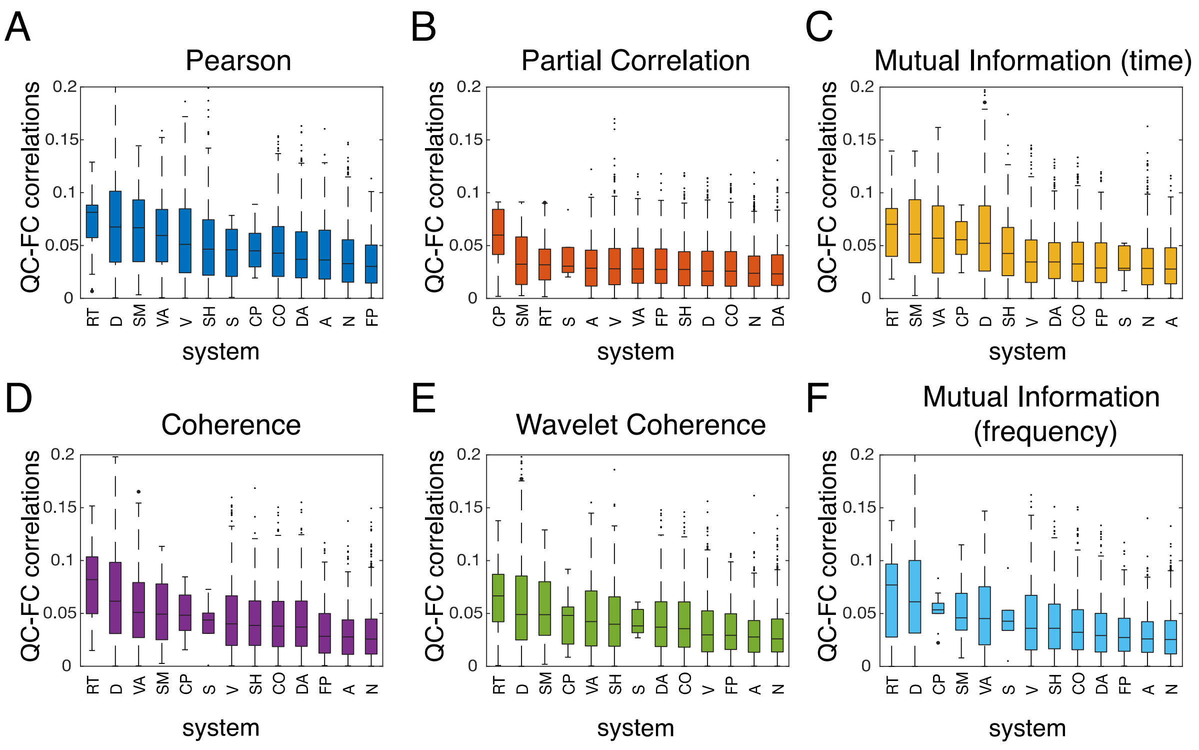


**Figure S4.** Boxplots of intra-system QC-FC correlations are shown for the REST1_LR scan and the 333-node Gordon parcellation, with descending median absolute QC-FC correlation from left to right. A=auditory; CO=cingulo-opercular; CP=cingulo-parietal; D=default; DA=dorsal attention; FP=fronto-parietal; N=none; RT=retrosplenial temporal; SH=sensory, motor, hand; SM=sensory, motor, mouth; S=salience, VA=ventral attention; V=visual.

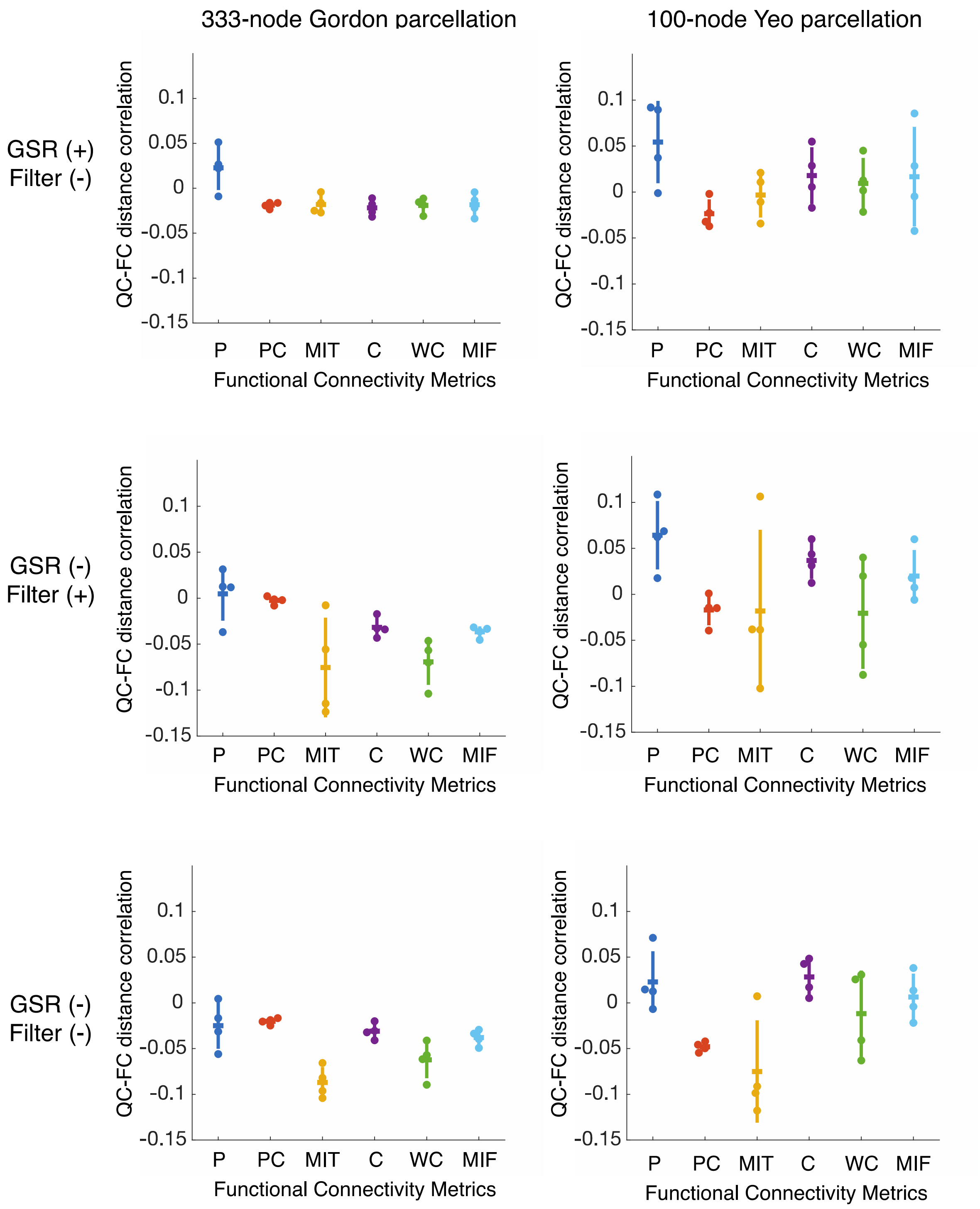


**Figure S5.** Residual distance-dependence of motion artifact for different FC estimation methods with three different preprocessing variants defined by the inclusion or exclusion of global signal regression (GSR) and bandpass filtering. **(A)** FC estimated using the 333-node parcellation provided by Gordon *et al.* (2016). **(B)** FC estimated using the 100-node parcellation provided by Schaefer *et al.* (2017); P=Pearson, PC=Partial Correlation, MIT=Mutual Information (time), C=Coherence, WC=Wavelet Coherence, MIF=Mutual Information (frequency). Notches represent the mean of four resting-state scans and error bars show the standard deviation.

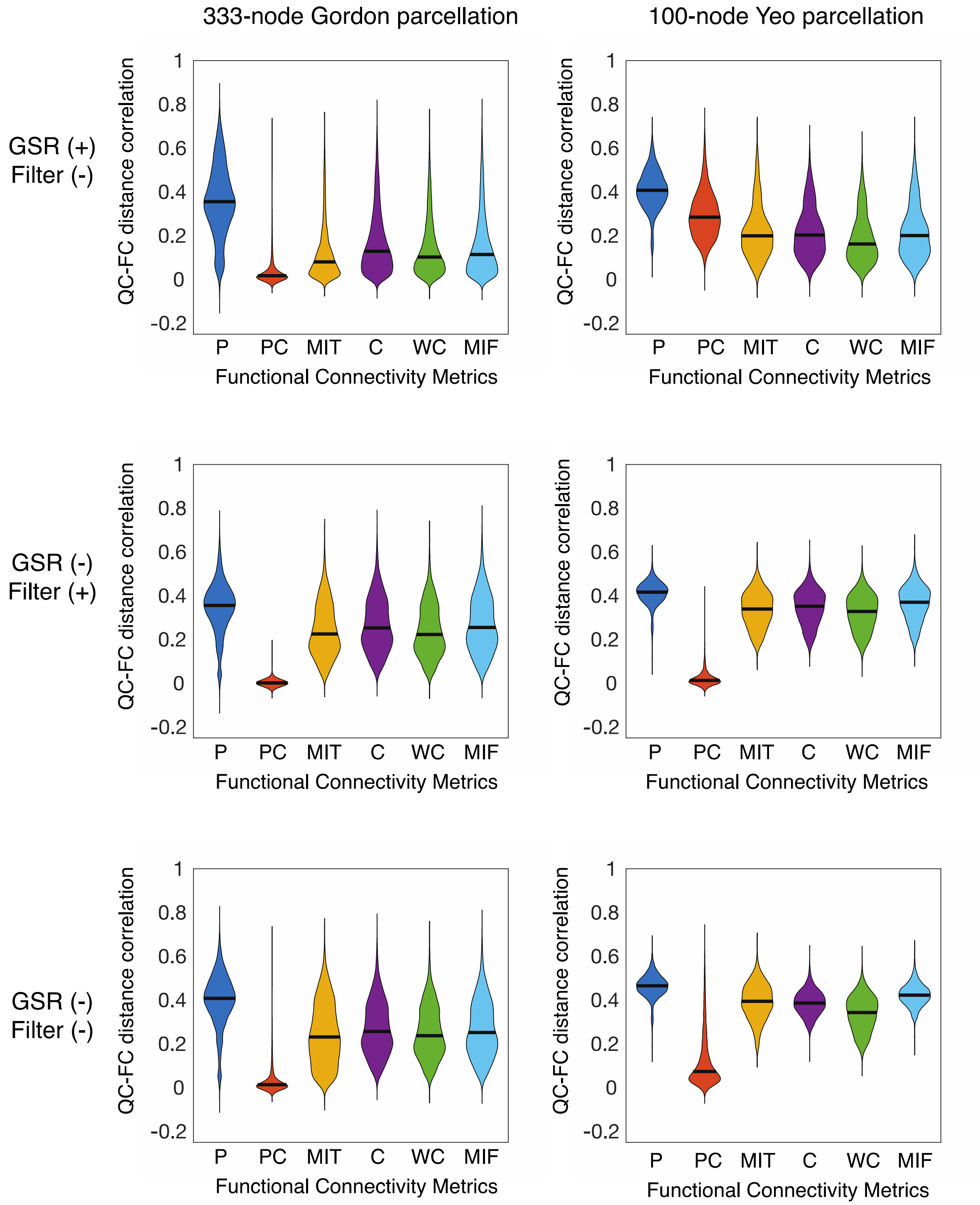


**Figure S6.** Intra-class correlation for all edge weights with three different preprocessing variants defined by the inclusion or exclusion of global signal regression (GSR) and bandpass filtering. **(A)** FC estimated using the 333-node parcellation provided by Gordon *et al.* (2016). **(B)** FC estimated using the 100-node parcellation provided by Schaefer *et al.* (2017); P=Pearson, PC=Partial Correlation, MIT=Mutual Information (time), C=Coherence, WC=Wavelet Coherence, MIF=Mutual Information (frequency). Notches represent the mean of four resting-state scans and error bars show the standard deviation.

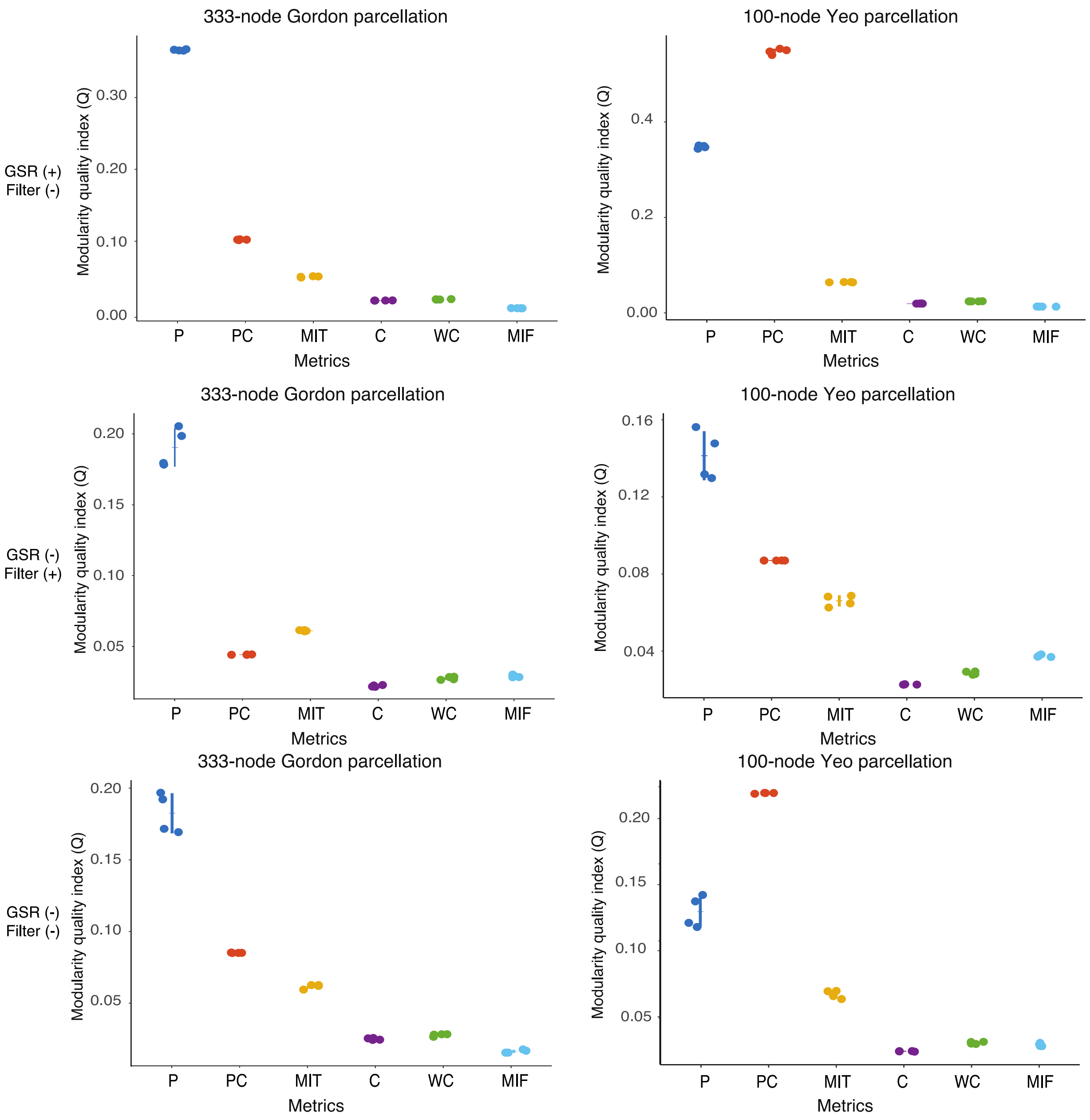


**Figure S7.** Modularity quality index with three different preprocessing variants defined by the inclusion or exclusion of global signal regression (GSR) and bandpass filtering. **(A)** FC estimated using the 333-node parcellation provided by Gordon *et al.* (2016). **(B)** FC estimated using the 100-node parcellation provided by Schaefer *et al.* (2017); P=Pearson, PC=Partial Correlation, MIT=Mutual Information (time), C=Coherence, WC=Wavelet Coherence, MIF=Mutual Information (frequency). Notches represent the mean of four resting-state scans and error bars show the standard deviation.

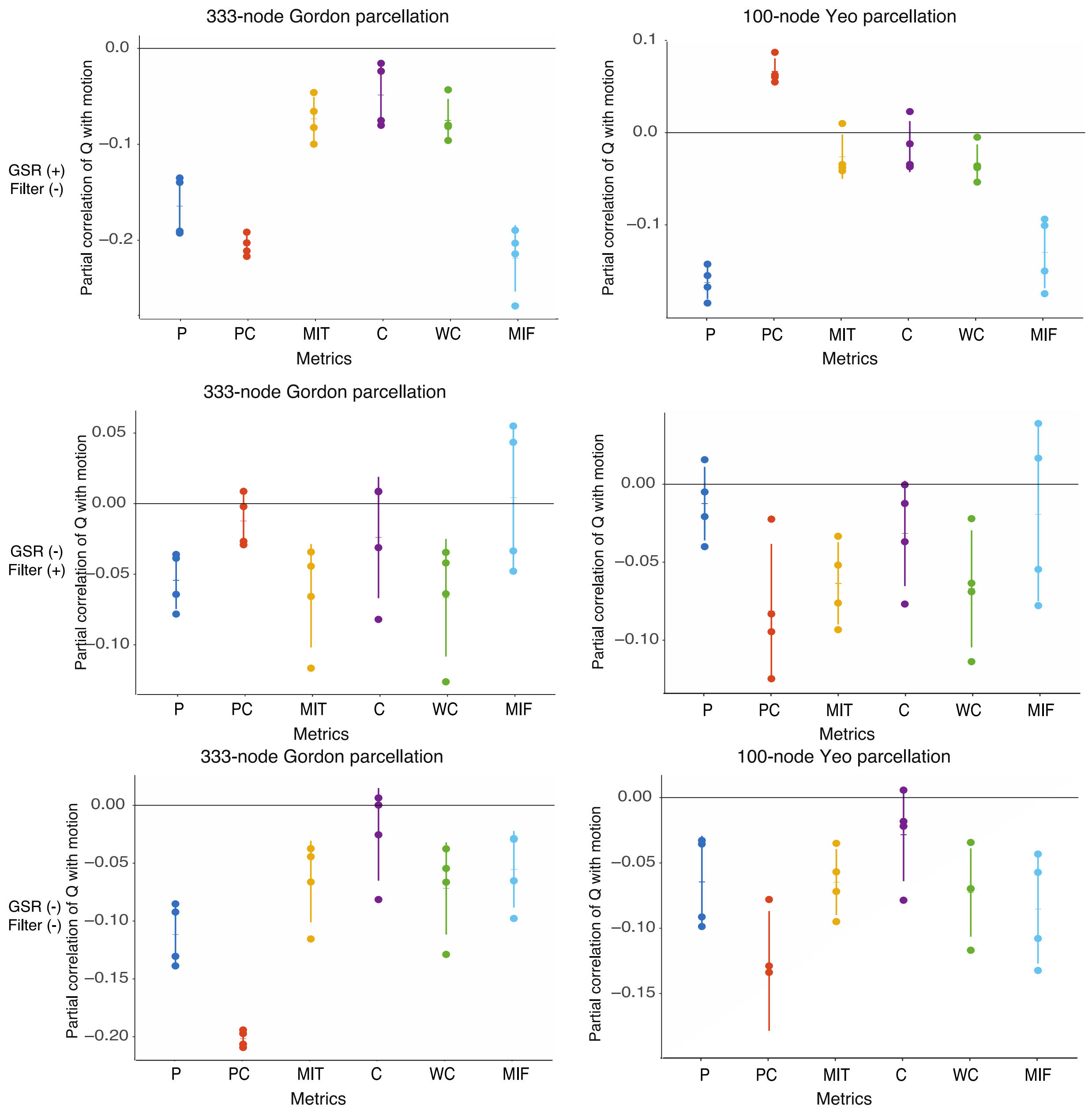


**Figure S8.** Partial correlation between $Q$ and mean relative RMS, controlling for average network weight, age, and sex, with three different preprocessing variants defined by the inclusion or exclusion of global signal regression (GSR) and bandpass filtering. **(A)** FC estimated using the 333-node parcellation provided by Gordon *et al.* (2016). **(B)** FC estimated using the 100-node parcellation provided by Schaefer *et al.* (2017); P=Pearson, PC=Partial Correlation, MIT=Mutual Information (time), C=Coherence, WC=Wavelet Coherence, MIF=Mutual Information (frequency). Notches represent the mean of four resting-state scans and error bars show the standard deviation.

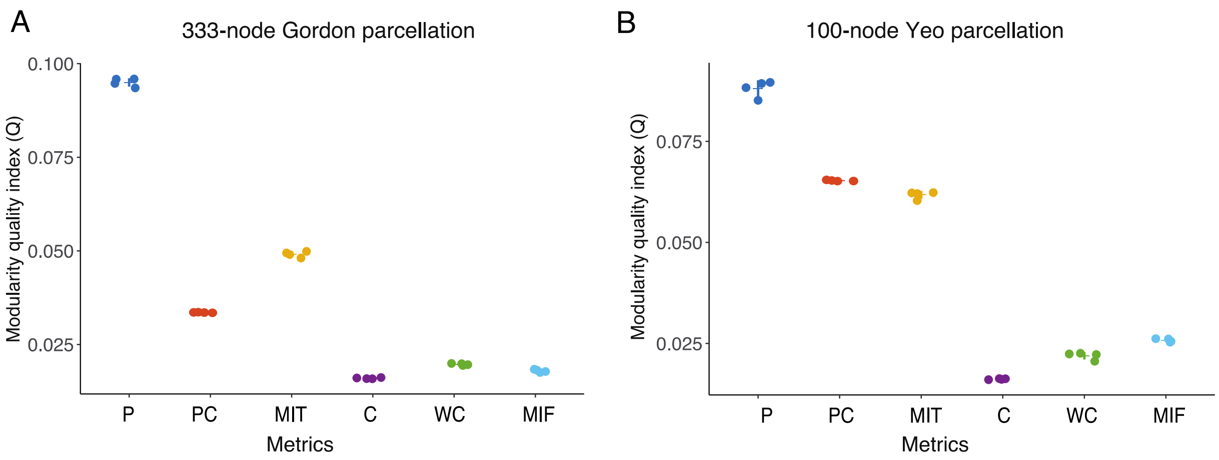


**Figure S9.** Modularity quality index estimated on networks containing only absolute values of edge weights. The weighted generalization of modularity maximization (Rubinov & Sporns, 2017) was used for all networks in this analysis. Modularity quality values are more comparable across metrics than when using signed matrices. **(A)** FC estimated using the 333-node parcellation provided by Gordon *et al.* (2016). **(B)** FC estimated using the 100-node parcellation provided by Schaefer *et al.* (2017). P=Pearson, PC=partial correlation, MIT=Mutual Information (time), C=Coherence, WC=Wavelet Coherence, MIF=Mutual Information (frequency).

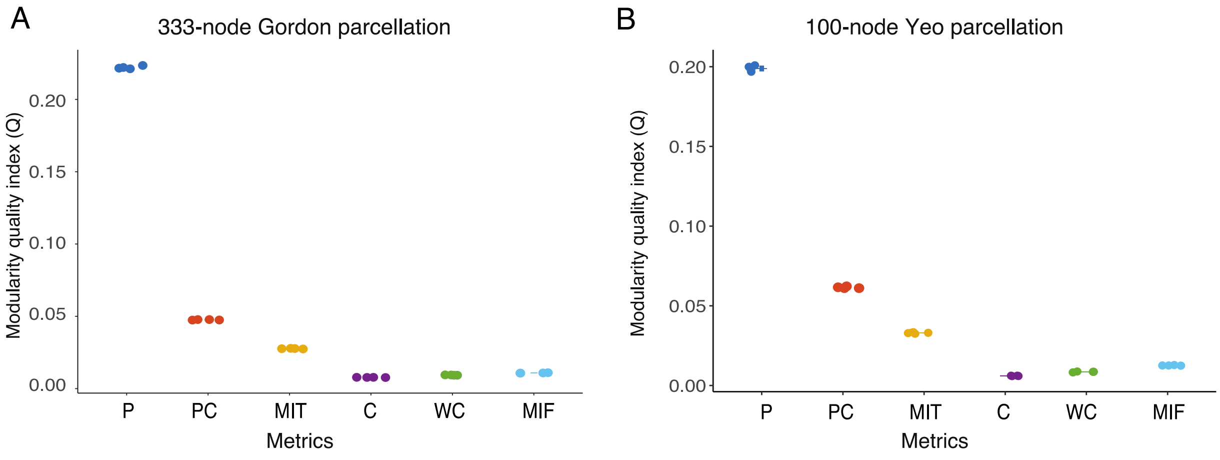


**Figure S10.** Modularity quality index estimated on networks using the *a priori* corresponding set of communities. For the Gordon parcellation, the accompanying 13-community partition was used. For the Schaefer 100-region parcellation, the 17-community partition was used. **(A)** FC estimated using the 333-node parcellation provided by Gordon *et al.* (2016). **(B)** FC estimated using the 100-node parcellation provided by Schaefer *et al.* (2017). P=Pearson, PC=partial correlation, MIT=Mutual Information (time), C=Coherence, WC=Wavelet Coherence, MIF=Mutual Information (frequency).

**Supplemental References**


1. Blondel, V. D., Guillaume, J. L., Lambiotte, R. & Lefebvre, E. Fast unfolding of communities in large networks. *J. Stat. Mech. Theory Exp.* **2008**, 0–12 (2008).
2. Ciric, R. *et al.* Benchmarking of participant-level confound regression strategies for the control of motion artifact in studies of functional connectivity. *Neuroimage* **154**, 174–187 (2017).
3. Thomas Yeo, B. T. *et al.* The organization of the human cerebral cortex estimated by intrinsic functional connectivity. *J. Neurophysiol.* **106**, 1125–1165 (2011).
4. van Wijk, B. C. M., Stam, C. J. & Daffertshofer, A. Comparing brain networks of different size and connectivity density using graph theory. *PLoS One* **5**, (2010).
5. Ginestet, C. E., Nichols, T. E., Bullmore, E. T. & Simmons, A. Brain network analysis: Separating cost from topology using Cost-Integration. *PLoS One* **6**, (2011).
6. Yan, C. G., Craddock, R. C., Zuo, X. N., Zang, Y. F. & Milham, M. P. Standardizing the intrinsic brain: Towards robust measurement of inter-individual variation in 1000 functional connectomes. *Neuroimage* **80**, 246–262 (2013).
7. Gordon, E. M. *et al.* Generation and Evaluation of a Cortical Area Parcellation from Resting-State Correlations. *Cereb. Cortex* **26**, 288–303 (2016).
8. Schaefer, A. *et al.* Local-Global Parcellation of the Human Cerebral Cortex from Intrinsic Functional Connectivity MRI. *Cereb. Cortex* **28**, 3095–3114 (2018).
